# Comprehensive analysis of the transcription factor REST regulatory networks in IDH-mutant and IDH-wild type glioma cells and gliomas

**DOI:** 10.1101/2022.07.28.501927

**Authors:** Malgorzata Perycz, Michal J. Dabrowski, Marta Jardanowska, Adria-Jaume Roura, Bartlomiej Gielniewski, Karolina Stepniak, Michał Dramiński, Bozena Kaminska, Bartosz Wojtas

**Affiliations:** Laboratory of Molecular Neurobiology, Nencki Institute of Experimental Biology, Polish Academy of Sciences; Computational Biology Group, Institute of Computer Science of the Polish Academy of Sciences; Doctoral School of Institute of Biochemistry and Biophysics, Polish Academy of Sciences; Laboratory of Sequencing, Nencki Institute of Experimental Biology, Polish Academy of Sciences

**Keywords:** differentiation, glioblastoma, IDH mutation, REST, KAISO, ZBTB33, DNA methylation, transcription factor, extracellular matrix, invasion

## Abstract

REST is a widely expressed, dual role transcription factor that acts either as a transcriptional repressor or transcriptional activator depending on the genomic and cellular context. REST is an important oncogenic factor, a key player in brain cell differentiation and has a role in establishing DNA methylation status in proximity of its binding sites. Mutations in IDH cause significant changes to the epigenome contributing to blocking cell differentiation and are considered an oncogenic driver in glioma. We aimed at defining the REST role in the IDH mutation-related phenotype in gliomas accounting for its role in gene activation and repression. We studied the effects of REST knockdown, REST binding sites, and REST motifs methylation in context of IDH mutation, and found that both REST binding patterns and TF motif composition proximal to REST binding sites differed in IDH wild-type and mutant glioma. Among such REST targets were genes involved in glial cell differentiation and ECM organization. REST knockdown differently impacted glioma cell invasion depending on the IDH phenotype. DNA methylation of REST activated gene promoters showed positive correlation with gene expression. The canonical REST-repressed gene targets correlated with NPC-like cellular state properties in IDH-MUT grade 2/3 gliomas. The identified REST targets, gene regulatory networks and putative REST cooperativity with other TFs point to differential control of REST target gene expression in IDH-WT and IDH-MUT gliomas. We conclude that REST could be considered as a key factor in the design of targeted glioma therapies.

## Introduction

REST (RE1-silencing transcription factor), previously known as NRSF – the neuron-restrictive silencer factor, participates in the control of neuronal differentiation, regulating the transition from stem to progenitor cells [1]. REST and its co-repressor complex participate in shaping neuronal plasticity in developing brains, as well as repressing neuronal gene expression in non-neuronal terminally differentiated cells [1]. REST is also one of the most prominent players in glioblastoma stem cells regulating their tumorigenic potential [2]. REST may recruit many epigenetic factors that in turn can repress or activate gene expression by imprinting active or repressive marks on histones and DNA [1,3–5]. REST can recruit MECP2 – methyl-CpG binding protein 2 and as a part of CoREST/MeCP2 repressor complex can bind to methylated sites distinct from canonical RE1 sites [1,4]. We have recently shown that DNA methylation within the predicted REST binding sites has a prognostic value in patients with higher grade gliomas [6]. Moreover, increased expression of genes putatively regulated by REST correlates with poor prognosis of glioma patients [7]. REST appears as a transcription factor which expression correlates well with its activity and its activity can be well inferred from the expression of its targets [7].

Mutations in genes coding for IDH1 or IDH2 (isocitrate dehydrogenases) occur frequently in oligodendroglioma WHO grade 2, 3 and astrocytoma grade 2, 3, 4 and result in global changes in DNA methylation. A mutated IDH enzyme (IDH-MUT, most often carrying a R132H substitution) catalyzes production of both, α -ketoglutarate (α-KG), and 2-hydroxyglutarate (2HG) [8]. 2HG oncometabolite can inhibit α-KG-dependent dioxygenases, such as DNA and histone demethylases [9]. As DNA and histone demethylation are inhibited, cells are blocked in the hypermethylated epigenetic state, which dysregulates expression of genes and blocks cell differentiation [10,11]. Whole genome epigenetic changes related to IDH1/2 mutations are often described as IDH-related phenotype [12].

REST can bind both unmethylated and methylated DNA in various contexts [2,13,14]. Knockdown of REST in glioblastoma (GBM) xenografts significantly impairs tumor growth [15]. The authors proposed that REST inhibits expression of neuronal lineage genes in glioma cells, leading to self-renewal and increasing tumorigenic potential of cells, while in the cells with REST knockdown, the inhibition of neuronal genes is abolished, leading to either cell differentiation or cell death [15]. The role of REST in gliomas is far from conclusive. On one hand, the inhibition of REST in established human U87 and U251 glioma cells resulted in reduced cell proliferation and migration [16]. On the other hand, REST and p53-deficient mice develop glioblastomas that are similar to human proneural tumors with some characteristics of primitive neuroepithelial tumors (PNET) [17]. Therefore, the role of REST varies depending on a cell origin and its developmental stage, suggesting it could be context dependent.

In the present study, we sought to determine at genome-wide perspective, whether inhibiting and activating functions of REST in glioma cells are modified by the IDH status and inferred changes in DNA methylation in gliomas and glioma cell lines.

## Materials and methods

### Cell culture

The experiments were conducted using U87-MG (ATCC HTB-14 IDH-WT) and IDH1 mutant-U87 Isogenic Cell Line (ATCC HTB-14IG), hereinafter referred to as U87-WT and U87-MUT, purchased from American Type Culture Collection (ATCC). The cells were cultured in DMEM (Gibco), 10% FBS (PAN-BIOTECH) with no antibiotics added. Cells were passaged every 3-5 days depending on the seeding density.

### Human glioma samples

This study used glioma tumor data generated in the work of Stepniak et al, 2021 [18] (EGA accession nr: ERP125425)

### REST-ChIP-seq in human glioma samples

Glioma tumors (n=7) REST-ChiP-seq results were obtained with a protocol described in Stepniak et al, 2021 [18] with the exception of using REST antibody (Millipore, CS200555) instead of a histone mark antibody.

### Human glioma samples: methylomes

5mC DNA methylation data (NimbleGen SeqCap Epi from ROCHE) include glioma tumor samples of WHO grade 2/3 IDH-WT (n=4), grade 4 IDH-WT (n=11) and four IDH-MUT samples (n=4), obtained from the recently published atlas of active enhancers and promoters in benign and malignant gliomas [18] and regulomics.mimuw.edu.pl/GliomaAtlas). In our study we refer to this dataset as “glioma Atlas”. All four IDH-MUT samples were pooled together and hereafter are referred to as IDH-MUT. The glioma Atlas data covers millions of cytosines at a base pair resolution. Due to tumor-related DNA degradation, methylomes presented in the Atlas consist of over 10 million of cytosines per sample on average. Despite the limited number of tumor type specific samples, the number of covered DNA sites in the Atlas is a huge advantage in comparison to BeadChip panels such as Infinium® HumanMethylation450 or MethylationEPIC performed on larger patient cohorts.

### Human glioma samples: RNA-seq

RNA-seq row counts of genes were obtained for the same set of samples from the glioma Atlas whose methylomes were analyzed (n=4 IDH-MUT, n=4 IDH-WT LGG (lower grade gliomas, WHO grade 2/3), n=11 IDH-WT GIV).

### TCGA public data

TCGA level 3 RNA-seq data repositories of GBM (glioblastoma, WHO grade 4) and LGG (lower grade gliomas, WHO grade 2/3), aligned by STAR and gene expression counted by HTseq, https://portal.gdc.cancer.gov/) were uploaded to R. Gene expression levels as FPKM (fragments per kilobase of exon per million) were used for further analysis of REST expression in the context of glioma malignancy grades and IDH1/2 mutation status. Clinical data for LGG and GBM datasets were obtained from [19] (Table S1 therein).

### Grade 4 primary glioma cell lines

RNAseq results (DESeq2 normalized) from 12 grade 4 glioma cell lines, 2 IDH-MUT astrocytomas and 10 GBM IDH-WT [20] were used for U87 IDH-MUT model validation in primary human glioma cell lines.

### Single cell RNA-seq data

Glioma tumor samples scRNA-seq data [21] were uploaded from GEO (GSE131928) and data were uploaded to R as TPM values.

### siRNA-REST knockdown

REST knockdown was performed in U87-WT and U87-MUT, the isogenic malignant glioma cell lines that genetically differed only by IDH1 mutation status. The cells were subcultured 2 days prior to the transfection so they would not exceed the confluency of 80% on the day of siRNA transfection and double transfected within 24 hours, first by nucleofection, followed by lipofection. For the nucleofection, cells were trypsinized, counted, centrifuged, resuspended in Lonza SE cell line solution reagent (Lonza, PBC1-02250) and transferred to Nucleocuvette Vessel (Lonza). Control or human REST-targeting siRNA ON-TARGETplus SMARTpool (Dharmacon, D-001810-10-05 and L-006466-00-0005) was then added to the appropriate wells of the vessel and nucleofection was carried out using 4D-Nucleofector core unit. The cells were then resuspended with DMEM 10% FBS and cultured in 12 or 24 well plates (Falcon) for the next 24 hours. After that time, the medium was replaced with fresh DMEM 10% FBS and the cells were transfected for the second time using Lipofectamine 2000 (Invitrogen, 2094065) and the same siRNA ON-TARGETplus SMARTpool (Dharmacon) as before. Protein and RNA were collected 72 hrs after the first transfection.

### RNA isolation

RNA was isolated from U87-WT and U87-MUT cells that were either not treated or transfected with control or REST-targeting siRNA. RNeasy Mini Kit (QIAGEN, 74106) was used according to the producer’s protocol. RNA concentration was measured with NanoDrop 2000 (Thermo Scientific, NanoDrop products, Wilmington, USA). RNA quality was verified with Bioanalyzer 2100 (Agilent Technologies, Santa Clara, CA) using an RNA 6000 Nano Kit (Agilent Technologies, Ltd., 5067-1511).

### Quantitative PCR (qRT-PCR)

Complementary DNA was synthesized from total RNA by extension of oligo(dT) primers with SuperScript III Reverse Transcriptase (Invitrogen, 18080-044). Real-time PCR was performed applying SYBR Green chemistry (Applied Biosystem by Thermo Fisher Scientific, 4385612) on QuantStudio 12 K Flex Real-Time PCR System. Amplified product was normalized to the endogenous expression of GAPDH and represented as -ΔΔCt (negative delta delta Ct) values (fold change). Statistical significance of comparisons between groups was calculated in GraphPad Prism v. 9.1.2 (GraphPad Software, LCC). P values were considered significant when *P < 0.05 (Mann-Whitney test). Data were obtained in 4 independent experiments.

### Western Blotting

Whole-cell protein extracts were prepared, resolved by electrophoresis, and transferred to a nitrocellulose membrane (GE Healthcare, 10600003) as described [22]. After blocking with 5% non-fat milk in TBST (Tris-buffered solution pH 7.6, 0.01% Tween-20) the membranes were incubated overnight with primary antibody (rabbit anti-REST1, Millipore CS200555, 1:1000, or mouse anti-GAPDH, Millipore MAB374, 1:500) in TBST with 5% bovine serum albumin (BSA, Sigma) or 1 h with horseradish peroxidase-conjugated anti-β-actin antibody (Sigma, A3854) diluted 1:20000 in 5% non-fat milk in TBST. The primary antibody reaction was followed by 1 h incubation with horseradish peroxidase-conjugated anti-rabbit IgG diluted at 1:10000 (Vector, PI-1000). Immunocomplexes were detected with an enhanced chemiluminescence detection system (ECL, BioRad) and Chemidoc (BioRad). The molecular weight of proteins was estimated with Cozy prestained protein ladder (High Qu GmbH, PRL0102c1). Densitometry of band intensities was performed using BioRad Image Lab software. REST band intensities were normalized to GAPDH band intensities for each blot. Statistical significance of comparisons between groups was calculated in GraphPad Prism v. 9.1.2 (GraphPad Software, LCC). P values were considered significant when *P < 0.05 (column statistics t-test). Data were obtained in 4 independent experiments.

### PrestoBlueTM Cell Viability assay

PrestoBlue™ Cell Viability reagent (Invitrogen, A13262) was used to assess cell viability. In this assay we used blank (no cells), not treated, mock transfected and siRNA-transfected U87-WT and U87-MUT cells. Transfection was carried out as described earlier. Cells were seeded at 24 well plates at 40k/well. Time points for measuring the viability were 12h, 24h, 48h and 72h post nucleofection. PrestoBlueTM Cell Viability reagent was added for 30 minutes incubation in 37℃ (with gentle shaking) at a final concentration of 1x. A portion of PrestoBlue diluted in the medium was kept at the same conditions to be used as a blank. After incubation, 100 ul of medium from each well was transferred to 96 well plate and the fluorescence was read at an excitation/emission wavelength of 560/590 nm using BioTek Synergy HTX fluorimeter. Data was analyzed in GraphPad Prism using Wilcoxon matched-pairs signed rank test, two-tailed. For each timepoint, mock transfected cells viability was normalized to 100%. Data were obtained in 3 independent experiments.

### Matrigel Invasion assay

Matrigel invasion assay was performed using tissue culture inserts (6.5 mm Transwell® with 8.0 μm Pore Polycarbonate Membrane Insert, Corning) coated with the Growth Factor Reduced Matrigel™ Matrix (BD Biosciences, 356231). 50 μL of the Matrigel™ Matrix (1 mg/mL) diluted in fresh DMEM medium was dried under sterile conditions (37°C) for 4.5-6 h. The medium was added to the wells 1 h before seeding the glioma cells into inserts. The U87-WT and U87-MUT cells were double transfected 54h prior to plating in matrigel-covered chambers; mock transfected and not transfected cells were used as control. The cells were seeded on matrigel-covered membrane at 45k per insert in 5% FBS-DMEM. The cultures were placed in a 37°C humidified incubator with 5% CO_2_. After 18h, the inserts were washed with PBS, had their inside cleaned with a cotton swab, and were placed in ice-cold methanol for 20 minutes to fix the cells that had migrated. The membranes were then cut out from the Transwell® inserts and mounted using VECTASHIELD Antifade Mounting Medium with DAPI (Vector Laboratories, H-1800). Cell images were taken within five independent fields (bottom, top, left, right side, and a center) of each specimen, using a fluorescence microscope (Leica DM4000B, 5× objective). Numbers of migrating cells’ nuclei were counted using ImageJ software. Experiments were performed in duplicates six times in total. Statistical analysis was performed using GraphPad Prism software. The groups were compared using Wilcoxon matched pairs signed rank test and differences considered significant when *P < 0.05.

### REST-Chromatin immunoprecipitation

U87-WT and U87-MUT cells were plated at 10 cm plates, and the following day the cells were trypsinized, centrifuged, resuspended in PBS and fixed with 1% formaldehyde for 10 minutes. The cells were lysed, and chromatin was isolated and fragmented by sonication (Covaris). Fragmented chromatin was loaded onto REST antibody (Millipore, CS200555) or IgG (Millipore, PP64B) - coated agarose beads (Invitrogen). Immunoprecipitated complexes were eluted, and DNA was purified using DNA Clean&Concentrator kit (Zymo Research).

### ChIP sequencing

DNA libraries for chromatin immunoprecipitation sequencing were prepared using QIAseq Ultra Low Input Library Kit (QIAGEN, ZZ-QG-180492) for two independent REST-ChIP experiments. Briefly, DNA was end-repaired, adenosines were added to the 3′ ends of dsDNA and adapters were ligated (adapters from NEB, Ipswich, MA, USA). Following the adapter ligation, uracil was digested by USER enzyme from NEB (Ipswich, MA, USA) in a loop structure of the adapter. Adapters containing DNA fragments were amplified by PCR using NEB starters (Ipswich MA, USA). Library quality evaluation was done with Agilent 2100 Bioanalyzer using the Agilent DNA High Sensitivity chip (Agilent Technologies Ltd, 5067-4626). Quantification and quality evaluation of obtained samples were done using Nanodrop spectrophotometer (Thermo Scientific, NanoDrop products), Quantus fluorometer and Quanti Fluor ONE dsDNA Kit (Promega, E4870) and 2100 Bioanalyzer (Agilent Technologies). The average length of the DNA fragments in the library was 300 bp. The libraries were run in the rapid run flow cell and were single end sequenced (65 bp) on HiSeq 1500 (Illumina).

### RNA sequencing

Total RNA was obtained from 3 passages of each U87-WT and U87-MUT that were either not treated, control- or REST-siRNA transfected as described earlier (Dharmacon, Lonza, and Invitrogen). 500 ng of RNA was used for cDNA synthesis for transcriptome sequencing. mRNA sequencing libraries were prepared using KAPA Stranded mRNAseq Kit (Roche, 07962193001), according to the manufacturer’s protocol. Briefly, mRNA molecules were enriched from 500 ng of total RNA using poly-T oligo-attached magnetic beads (Kapa Biosystems). The first-strand cDNA was synthesized using a reverse transcriptase. Second cDNA synthesis was performed to generate double-stranded cDNA (dsDNA). Then, the adapter was ligated, and the loop structure of the adapter was cut by USER enzyme (NEB, Ipswich, MA, USA). Amplification of obtained dsDNA fragments containing the adapters was performed using NEB starters (Ipswich, MA, USA). Quality control of obtained libraries was done using Agilent Bioanalyzer with High Sensitivity DNA Kit (Agilent Technologies, Palo Alto, CA, USA, 5067-4626). Quantification of the libraries was done using Quantus Fluorometer and QuantiFluor Double Stranded DNA System (Promega, Madison, Wisconsin, USA, E4870). The libraries were run in the rapid run flow cell and were paired end sequenced (2×76bp) on HiSeq 1500 (Illumina, San Diego, CA 92122 USA).

### Glioma Atlas DNA methylation

DNA raw methylation data were provided by the Authors of the glioma Atlas. The data were processed with CytoMeth tool to obtain DNA methylation level as beta value (β value) in a single nucleotide resolution.

### Whole genome DNA methylation of U87 cell line samples

Twelve U87-MG cell line samples were analyzed in total: six IDH-MUT samples, out of which three were siREST and three siCTRL treated. The other six samples were IDH-WT, out of which three were siREST treated and three siCTRL.

Briefly, genomic DNA was first mechanically fragmented using a Covaris M220 instrument to produce DNA fragments of 350-400 bp. The next steps included repair of the ends of the obtained fragments, adapter ligation, enzymatic conversion of cytosines and amplification of the library using PCR. Basic quality parameters of ready libraries were checked using Agilent Bioanalyzer (fragment length distribution) and Promega Quantus (concentration) devices. They were found to be correct for all samples. The average length of the library was 482 bp (SD=11), which is within the range defined as an appropriate (470-520 bp) by the kit manufacturer. Next-generation sequencing libraries were prepared using the NEBNext® Enzymatic Methyl-seq Kit (New England Biolabs, cat. no. E7120). Libraries were sequenced using a NovaSeq 6000 device in paired-end sequencing mode 2x150 cycles.

Complete methylomes of IDH-MUT (siREST, siCTRL) and IDH-WT (siREST, siCTRL) were intersected with all the REST-ChIPseq peaks (n=3833). Sites with a coverage of less than 7 reads were excluded from further analyses. A mean DNA methylation beta value for each cytosine locus was calculated for each sample type (WT-siCTRL, WT-siREST, MUT-siCTRL, MUT-siREST). Then the difference in DNA methylation level as well as fold change was computed between certain groups of samples. The difference in DNA methylation beta value >=0.15 was assumed as significant. Chi2 test was used to compare the distribution of background cytosines (all cytosines within REST-peaks) with differentially methylated cytosines (showing higher or lower methylation pattern in IDH-MUT cell lines) across common, IDH-WT or IDH-MUT specific REST-ChIPseq peaks, results were found significant with p<0.001.

### Glioma Atlas RNA-seq data

RNA-seq raw data were provided by the Authors of the glioma Atlas. Following the recommendations, two normalization procedures were considered: (i) within-lane to adjust for GC-content and gene-length; (ii) between-lane, both implemented in EDASeq 3.8 R package [23]. The normalized gene expressions were used to investigate correlation of DNA methylation level with gene expression.

### DiffMeth module for DNA methylation analysis

DiffMeth is a module that identifies statistically significant differences in methylation level of the given DNA regions among defined groups of samples. The input provided to this automatic analysis consisted of: (1) DNA regions of interest defined in a standard bed file (chromosome, starting position, ending position, name of the region, name of the gene, strand); (2) A set of standard bed files resulting from CytoMeth processing, containing cytosine methylation levels expressed as beta values for specific cytosine position on a given chromosome for a given sample. Each sample was defined by a separate bed file; (3) A csv file defining the sample set, which contained methylation file name, sample name/ID, sample group (eg various types of cancer to be compared), ignore flag, and (4) Diffmeth input parameters defined in yaml file. The first step of the analysis (significance criterion) is based on a standard chi2 statistical test where all groups are compared to each other (pair by pair) to discover possible differences between two groups out of *n* provided. Chi2 test compares distribution of beta values that belong to one of the following ranges: [0.0-0.2], [0.2-0.6], [0.6-1.0] for a given region, two compared groups and CG context. Finally in this step p-value is calculated (corrected by FDR). The second step (significance criterion) is based on Kruskal-Wallis statistical test applied on all groups at once similarly as before: for a given region and CG context and p-values corrected by FDR. The final interesting differential regions can be selected by intersection of results from both criteria where p-value<0.05.

### Patient survival analyses

TCGA transcriptomic data (from both the LGG and GBM TCGA transcriptome datasets) were used to conduct four distinct survival studies, in which the patients were stratified into two subgroups based on REST expression level (high REST mRNA and low REST mRNA): the first one considered GBM patients with IDH-WT phenotype; the second one considered separate cohorts of GBMs and LGGs and (WHO 2/3) patients harboring IDH1/2 mutations; the third one included patients with all glioma grades; and the fourth one that included LGG patients with wild-type IDH genes. Survival and survminer R libraries were used in the survival analyses and censored patients were included. To support the association between REST expression levels and patient survival, Kaplan-Meier estimators and the log-rank test were calculated.

### U87 cell line IDH-phenotype cross validation with TCGA glioma data

In order to validate the U87 IDH-phenotype model, gene expression differences between U87-WT and its isogenic U87-MUT cell line were compared to the IDH-phenotype in the TCGA glioma transcriptomes dataset. First, log2 fold change (log2 FC) values of differentially expressed genes (DEGs) between U87-WT and U87-MUT were compared to log2 FC values from TCGA LGG IDH-MUT (defined in TCGA as a mutation in *IDH1/2*) vs IDH-WT comparison. To validate the relevance of overlap of DEGs with the same direction of change in U87 and TCGA tumors MUT-WT comparisons, a bootstrapping technique (sampling 10000 times) was applied. Next, tumor samples from the TCGA repository (WHO grades 2-4, GBM + LGG set) were divided into IDH-WT (n=86) and IDH-MUT (n=365). Two differential analyses were carried out: 1) LGG IDH-WT vs LGG IDH-MUT and U87 IDH-WT vs U87 IDH-MUT. REACTOME Pathways enrichment analysis was performed on a selection of the most significantly changed DEGs (FDR<0.01) obtained from differentially up-(log2 FC>0) and down-regulated (log FC <0) genes, separately.

### Transcriptomic data

RNA sequencing reads were aligned to the human genome with the STAR algorithm [24], a fast gap-aware mapper. Then, gene counts were obtained by featurecounts [25] using human genome annotations. The counts were then imported to R and processed by DESeq2 [26]. The counts were normalized for gene length and library size and statistical analysis was done by DESeq2 for the following comparisons: IDH-MUT/WT isogenic cell lines, siREST/siCTRL treated IDH-MUT cell line and siREST/siCTRL treated IDH-WT cell line. Gene pathways analysis (KEGG, Gene Ontology, REACTOME) were performed using clusterProfiler R library [27]. Gene Ontology Biological Processes (GO BP) analysis for DEGs arising from REST knockdown in U87-WT and U87-MUT was done separately for down-(log2 FC <0 and adjusted p-value <0.05) and up-regulated (log2 FC >0 and adjusted p-value <0.05) genes from a pool of IDH-MUT and IDH-WT DEGs.

### REST ChIP-seq data processing

Template amplification and cluster generation were performed using the TruSeq SBS Kit v3 and 50 nucleotides were sequenced with Illumina HiSeq 1500. After quality filtering (average Phred >30) and removal of duplicates, reads were mapped to the human genome (hg38) with the BWA MEM tool. Samtools view was used to filter best quality reads (-q 20 and -F 256 flags) before peak calling. The peaks were called using the Model-based Analysis of ChIP-Seq (MACS2) algorithm (Feng et al. 2012) with default parameters; peaks with q-value <0.01 were considered in the analysis. Peaks were annotated to genes using ChIPseeker [28] and subsequent pathway analysis was performed with clusterProfiler [27]. The raw ChIP-seq data were deposited in the NCBI GEO database: GSE174308. For further analysis, the consensus REST ChIP-seq peaks were generated as follows: the datasets for experiment repeat 1 and 2 were intersected for each U87-WT and U87-MUT datasets, separately. The resulting peaks (present in both repeats of the experiment) were then limited to the length of 200 bp (±100 bp from the peak summit, that was defined as a middle point of the obtained peak) for further TF motifs and DNA methylation analysis. Peaks were centered and limited to 200 bp in order to have a more consistent and homogenous set of observations. The resulting consensus peaks for U87-WT and U87-MUT were then intersected again to yield peaks common to both U87-WT and U87-MUT, peaks were also subtracted from each other to receive peaks specific to U87-WT and specific to U87-MUT.

### Defining REST-activated and REST-repressed genes

The putative REST target genes were defined as those with a REST-ChIP-seq peak within a promoter [28] in at least one of the ChIP-seq experiment repeats in either U87-WT or U87-MUT. Annotation of REST-ChIP-seq peaks to promoter genes was performed with ChIPseeker. REST-ChIP-seq peak U87-WT and U87-MUT datasets for both sets were pooled. The peak summits were assigned as the middle point of each existing peak, and 200 bp peak sequences were generated covering 100 bp upstream and downstream of the summit. Next, genes defined as REST targets by ChIPseeker based on U87-WT and U87-MUT ChIP-seq experiments were correlated (Pearson’s correlation) with REST expression levels in the joined LGG/GBM TCGA transcriptome dataset. Genes that had significant positive correlation (correlation coefficient >0.15, Bonferroni corrected p-value <0.0001) of their mRNA level with that of REST were defined as REST-activated, while these having significant inverse correlation (correlation coefficient < -0.15, Bonferroni corrected p-value <0.0001) were defined as REST-repressed.

### Intersection of REST knockdown and REST ChIP-seq data

Gene targets of REST defined by REST ChIP-seq (588 activated, 981 repressed) were intersected with REST gene targets as defined by REST knockdown. The REST ChIP-seq gene targets that had decreased expression upon REST knockdown were defined as REST activated targets, while those that had increased expression upon REST silencing were defined as REST repressed targets. Separately, Gene Ontology Biological Process (GO BP) analysis was performed for both groups using clusterProfiler and GO BP database.

### Intersection of REST ChIP-seq peaks with ENCODE database

REST ChIP-seq peaks (peak summit ± 100bp) were submitted to the Enrichr tool (https://maayanlab.cloud/Enrichr/) to predict transcription factors binding sites deposited in ENCODE.

### A position weight matrix (PWM) analysis

To identify the transcription factors (TF) motifs within the REST ChIP-seq peaks a PWM analysis was performed on two datasets. The first dataset consisted of the sequences assigned to REST-activated (n=588) and REST-repressed genes (n=981). The second dataset contained sequences stratified according to the IDH gene mutation status (WT-specific n=1007, MUT-specific n=114, peaks common between both n=2647). The 200 bp peak sequences were used as an input for known motif search using the PWMEnrich Bioconductor R package [R-PWMEnrich] and open-access HOCOMOCO [29] database with 14 additional REST matrices obtained from ENCODE. To identify TF motifs, 10765 selected promoters active in tumor samples were used as a background (dataset derived from [18]) to proceed lognormal distribution background correction. Only these motifs for which the p-value of motif enrichment was lower than 0.001 were selected for further analysis. Due to the notable difference in the number of sequences between IDH-WT (n = 1007) and IDH-MUT (n = 114), ten draws of 114 sequences from the IDH-WT sequence pool were made. A separate analysis of the search for motifs using the PWMEnrich R package was performed on each of these ten sets. In parallel, the same analysis was performed on the set of all the sequences characteristic for IDH-WT (n = 1007). For further IDH status analysis, motifs common between the sum of TF motifs from ten draws and the motifs identified on the set of all available sequences were selected. To identify transcription factor binding sites (TFBS), defined as each occurrence of the motif in the sequence, an online FIMO tool [30] from MEME Suite 5.0.1 was used. FIMO scans sequences as double stranded and matches motifs to both forward and backward strands. The p-value threshold for the FIMO output was set to 0.001. For further analysis, only the TFBS with the q-value lower than 0.05 were considered (correction done using Benjamini and Hochberg procedure). The TF motifs specific for these TFBS were grouped by sequence using the STAMP website tool (http://www.benoslab.pitt.edu/stamp/). To compare motif sequences during grouping, the Pearson correlation coefficient was used. As an aligning method, the ungapped Smith-Waterman followed by an iterative refinement multiple alignment strategy was used. Finally, the UPGMA tree-building algorithm was used. An enrichment analysis was done using Fisher’s exact test with an FDR correction. Transcription factor families were identified based on the HOCOMOCO database, which uses data from the Classification of Transcription Factors in Mammalia database (https://hocomoco11.autosome.ru/human/mono?full=true). The CentriMo tool from MEME Suite 5.4.1 (https://meme-suite.org/meme/tools/centrimo) allowed to determine the position distribution of each motif on the sequences, check the local enrichment of the given sequences with selected motifs, and calculate the static significance of the result using the binomial test. The analysis was performed for the entire sequence length assuming E-value threshold ≤ 10.

### REST peaks hierarchical clustering

To cluster by similarity the peaks with detected REST and KAISO motifs, each peak was changed into a binary vector, where a REST or KAISO motif was assigned value=1 if present in a peak or value=0 if absent in a peak. Such vectors were used as the input into hierarchical clustering performed with the Heatmap function from the ComplexHeatmap R package, using the default setting. Grouping was done by rows.

### Glioma Atlas DNA methylation data analysis

Discretization of β-value: DNA methylation β-value ranging from 0 to 1 was discretized into three classes: low/hypomethylation [0-0.2], medium (0.2-0.6] and high/hypermethylation (0.6-1].

Selecting regions with differential DNA methylation: DNA methylation analysis was performed on the two region types: peaks that cover a sequence of ±100 bp from the peak summit, and TF site motifs that mean a range of DNA sequence where the presence of a TF motif was statistically confirmed. We used the DiffMeth module with false discovery rate correction (FDR) for multiple testing, to identify regions with significantly different DNA methylation patterns. The results with FDR<0.05 were considered statistically significant. In the context of TF motifs differential analysis, where the motifs frequently overlap with each other, a repeated selection of the same cytosine for a single TF data set was prevented. If a cytosine was located within multiple motifs of a single TF due to the motifs overlapping, its beta value was counted only once. When a cytosine was within motifs of different TFs (REST or KAISO), its beta value (β) was counted separately for motifs of a TF.

Comparison of DNA methylation between classes: To compare the number of low, medium, and high-methylated cytosines between the classes we used the Chi-Square test. To compare DNA methylation (β-value) between two classes we used the Wilcoxon rank sum test, and for more than two classes we used the Kruskal-Wallis test. The results with p<0.05 were considered significant.

### Correlation of gene expression with glioma cell states

Data were analyzed in R, expression of investigated gene was correlated (Pearson’s correlation) with certain cell states: NPC-like1, NPC-like2, MES-like1, MES-like2, OPC-like, AC-like, G1S and G2M as defined by [21].

## Results

### *REST* expression is positively correlated with glioma malignancy and IDH mutation

We analyzed *REST* expression in the Cancer Genome Atlas (TCGA) transcriptomic data encompassing normal brain (NB) tissues, LGG (Lower Grade Gliomas, WHO grades 2 and 3) and GBM (glioblastoma, WHO grade 4) samples. The expression of *REST* was the lowest in normal brain tissue and the highest in the WHO grade 4 gliomas (G4), with *REST* expression increasing with grade malignancy from G2, through G3 toG4 (Figure 1A). The differences in *REST* expression in NB, G2, G3 and G4 were significant (adjusted p-value < 0.001). Interestingly, within the LGGs, *REST* expression was significantly higher in *IDH1/2* mutated (IDH-MUT) gliomas than in wild type samples (IDH-WT) (Figure 1B; two-sample Wilcoxon test).

**Figure 1.**
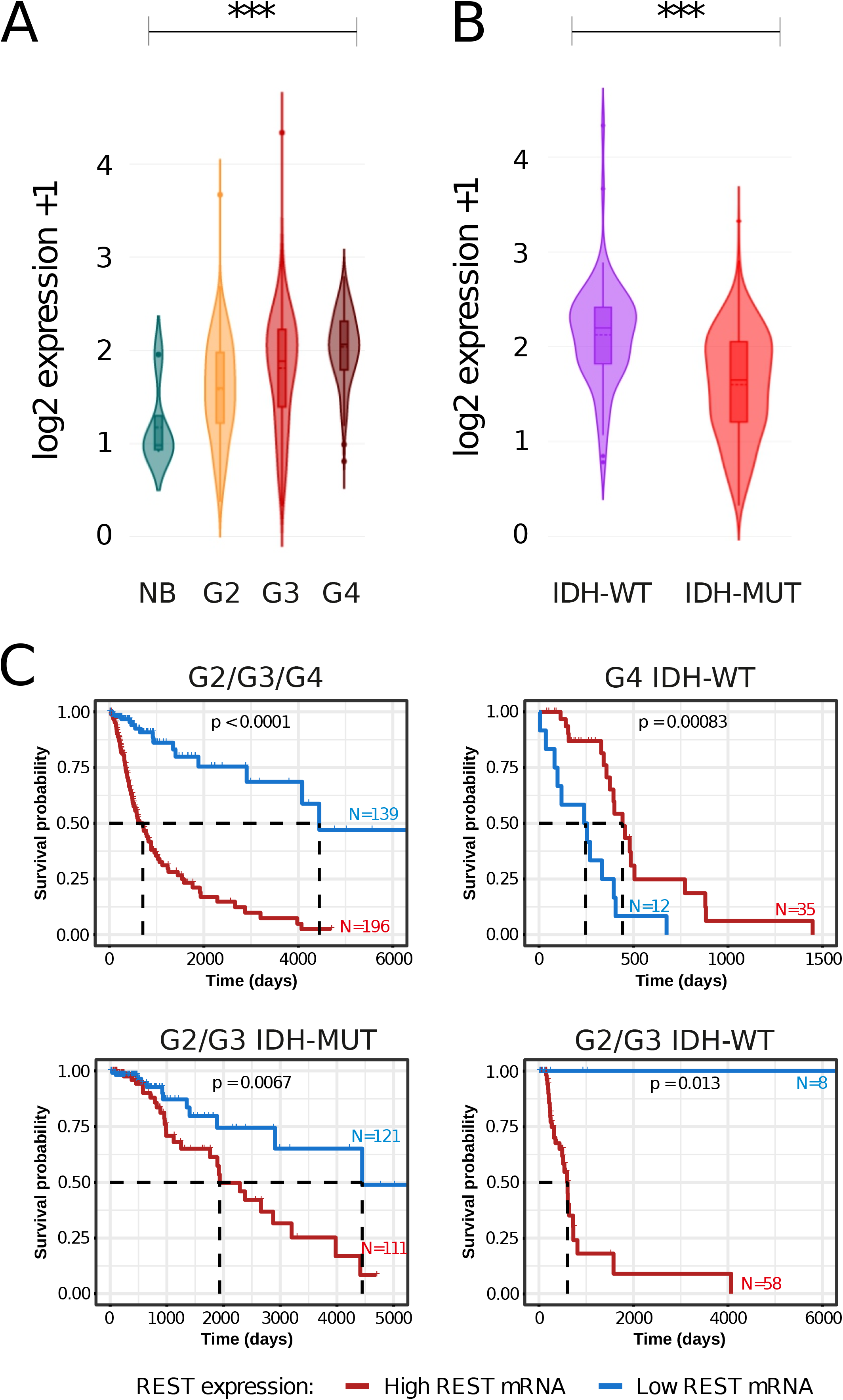
*REST* expression and survival analysis in the glioma TCGA dataset. **(A)** Violin plots showing expression of *REST* in a TCGA dataset across different glioma grades and normal brain. Significance of difference in gene expression was calculated with the Welch’s ANOVA test. ***=p-value<0.001. **(B)** Violin plot showing *REST* expression between *IDH1* wild type and *IDH1* mutants in G2/G3 gliomas (statistical significance calculated with the two-sample Wilcoxon test). ***=p-value<0.001. **(C)** Kaplan-Meier overall survival curves for the patients with WHO grade 2, 3, and 4 glioma (top left); IDH-WT G4 glioma (top right); IDH-MUT G2/G3 glioma (bottom left); grade 2 and 3 IDH wild-type (bottom right). The patients were divided into high or low *REST* expression groups. The patients alive at the time of the analysis were censored at the time of the last follow-up. Statistical significance was computed using the Log Rank Test.

Next, we investigated whether *REST* expression had a prognostic value in glioma patients’ survival. We confirmed that *REST* expression level is a strong negative prognostic factor for patient’s survival in the joint cohort of LGG and G4 samples. Patients with low *REST* expression had a favorable prognosis (Figure 1C, top left). When analyzed separately, patients with IDH-WT GBMs and high *REST* expression had longer survival (Figure 1C top right), contrary to patients with IDH-MUT LGGs (Figure 1C, bottom left) and IDH-WT LGGs (Figure 1C, bottom right), where high *REST* expression was associated with shorter survival. However, the results of IDH-WT LGG must be taken with caution since the sample size was small due to a scarcity of those tumors.

### Differentially expressed genes in human IDH1-WT and IDH1-MUT U87 cells and the TCGA dataset

We reasoned that before performing experiments on the U87 cell lines we will investigate their transcriptomes and compare them to tumor samples to validate this model. We took advantage of having isogenic U87 glioma cells with different *IDH1* status and analyzed transcriptomic profiles of U87 IDH-WT and U87 IDH-MUT glioma cells. We found significant differences in gene expression confirming transcriptomic deregulation (Additional File 1A). The REACTOME pathway analysis of differentially expressed genes (DEGs) in IDH-WT and IDH-MUT cells revealed a vast number of genes associated with extracellular matrix (ECM) and its reorganization (Additional File 1B). Interestingly, when DEGs identified in U87 cells were compared with the TCGA dataset a set of genes downregulated in U87 IDH-MUT as compared to U87 IDH-WT was highly concordant with a set of genes downregulated in IDH-MUT versus IDH-WT gliomas from TCGA (61% concordance, Figure 1A). Only TCGA LGG samples were included into this comparison because a number of IDH-MUT G4 samples was insufficient. Application of the bootstrapping method with 10,000 sampling ensured that such concordance is unlikely to be random as none of the bootstrapped results returned such a high concordance. The overlap of upregulated genes between glioma cell lines and TCGA gliomas was smaller (38%), which was less than the median overlap (48%) returned by bootstrapping. Genes upregulated in IDH-MUT U87 cells were enriched in REACTOME pathways that were dissimilar to those detected in LGGs (Additional File 1C). The REACTOME pathways of downregulated genes in U87 IDH-MUT vs U87 IDH-WT were highly similar to those from IDH-MUT vs IDH-WT in LGGs (Additional File 1D). Altogether, patterns of downregulated genes in IDH-MUT U87 glioma cells resulting from the hypermethylator phenotype well reflect changes detected also in IDH-MUT LGGs, while in case of upregulated genes the concordance is moderate.

### REST knockdown in U87 glioma cells affects numerous biological processes

To find REST dependent genes in glioma cells with different IDH1 status we performed siRNA mediated knockdown of REST (siREST) in IDH-WT and IDH-MUT U87 cells. A significant reduction in REST mRNA (Figure 2B) and protein (Figure 2C) levels was observed after 72 hours of REST silencing in both U87 cell lines. REST mRNA levels were reduced by 77% in IDH-MUT, and by 82% in IDH-WT (Figure 2B), while REST protein levels were reduced by 89% in IDH-MUT and by 90% in IDH-WT as compared to control siRNA (siCTRL) transfected cells (Figure 2C).

**Figure 2.**
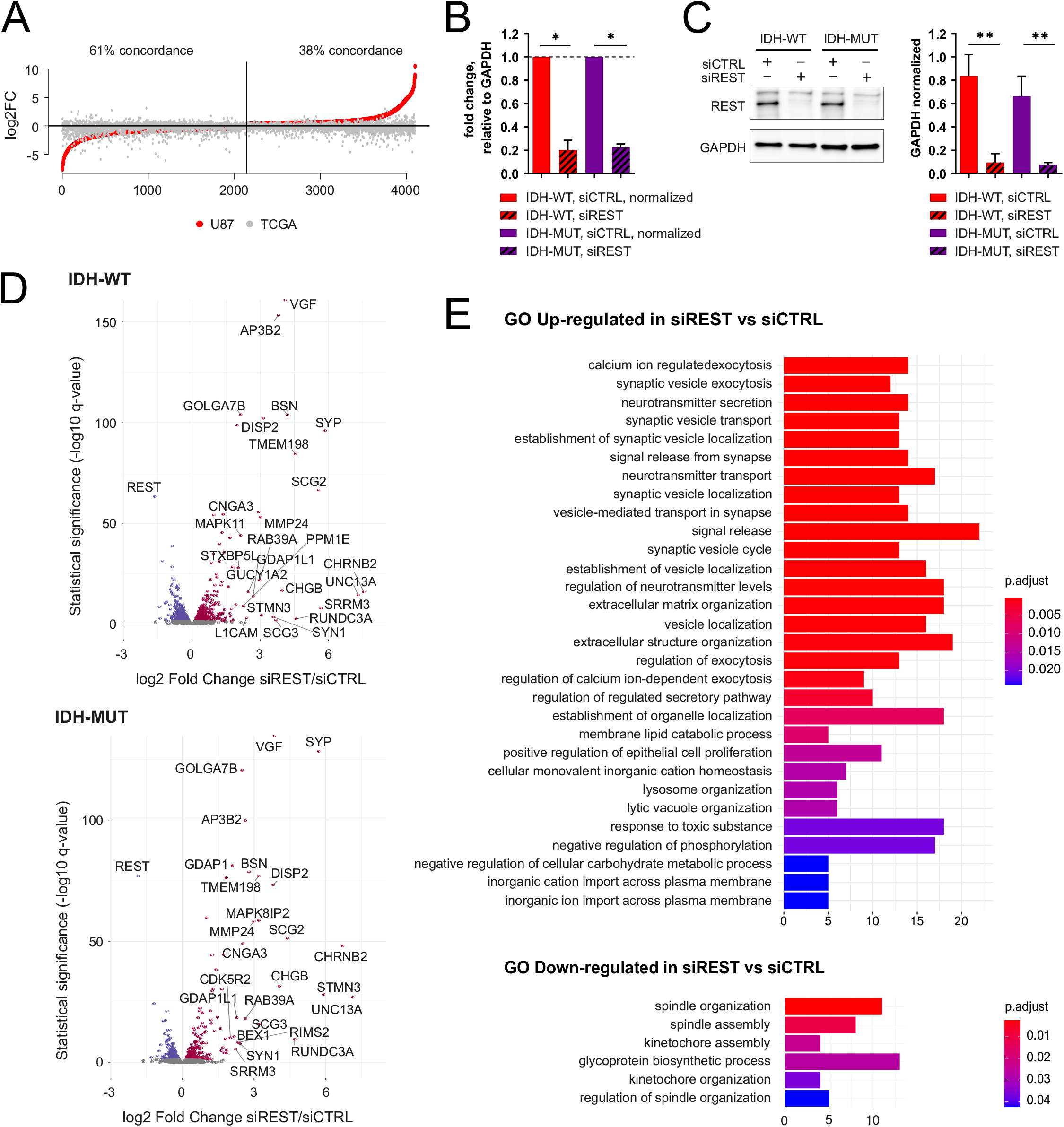
Effects of REST knockdown on gene expression in IDH-WT and IDH-MUT U87 cells. **(A)** Sorted values of log2 fold change (log2 FC) for the genes coming from the IDH-MUT vs IDH-WT comparison in U87 cell lines were presented as red dots. Values for the same genes coming from IDH-MUT vs IDH-WT comparison in TCGA G2/G3 gliomas were overlaid as gray dots. Percent of log2 FC direction concordance between IDH-MUT vs IDH-WT in U87 cell lines and glioma tumors was calculated. Number of differentially expressed genes is indicated. Black vertical line separates genes expressed higher in IDH-MUT U87 compared to IDH-WT U87 from the genes expressed higher in IDH-WT U87 compared to IDH-MUT U87. **(B)** Relative expression of REST in IDH-WT and IDH-MUT U87 cells at 72 hours of REST silencing with siRNA. mRNA levels in transfected cells were determined with quantitative PCR and normalized to GAPDH expression in the same sample. Data are represented as mean ± SEM, n=4 independent experiments, * p<0.05, two-tailed Mann-Whitney test (WT: p=0.0286; MUT: p=0.0286). **(C)** Levels of REST protein in IDH-WT and IDH-MUT U87 cells at 72 hours after transfection with control or REST specific siRNAs determined with Western blotting. Immunoblots were analyzed densitometrically. Data are represented as mean ± SEM, n=4. **p<0.01 (WT: p=0.0018, MUT: p=0.0052; two-tailed ratio-paired t-test). No difference was observed in the level of REST protein between silencing controls (siCTRL) in IDH-MUT and IDH-WT (p=0.226). **(D)** Volcano plots of the genes differentially expressed between siCTRL and siREST-transfected IDH-WT (upper plot) or IDH-MUT (bottom plot) U87 glioma cells. The axes show log2 fold change (x-axis) and -log10 from adjusted q-value (y-axis). **(E)** Gene Ontology (GO) Biological Processes (BP) analysis was performed on DEGs common in IDH-WT and IDH-MUT U87 cells. The results are presented as bar plots for pathways downregulated (upper panel) and upregulated (bottom panel) in REST depleted cells.

We compared transcriptomic profiles between siREST and siCTRL cells and identified 1,356 DEGs in IDH-WT cells and 629 in IDH-MUT (Figure 2D). Out of 1,356 DEGs, 507 were common for IDH-WT and IDH-MUT cells. The majority of the common DEGs were upregulated (n=287) after REST knockdown, whereas 220 DEGs were downregulated, including *REST,* which had the highest log2 fold change (log2 FC) between siREST and siCTRL in both IDH-WT and IDH-MUT models (Figure 2C, D). The Gene Ontology Biological Processes (GO BP) functional analysis was performed independently for the siREST up- and downregulated DEGs. The downregulated DEGs were enriched in cell division-related pathways (Figure 2E, upper panel), whereas the upregulated genes were enriched in neuronal-specific pathways, as well as pathways associated to endothelial cell proliferation and extracellular matrix organization (Figure 2E, bottom panel). However, analysis of IDH-WT vs IDH-MUT in both U87 cell lines and tumors revealed that ECM-related pathways may be differentially regulated, pointing to REST as a potential modulator of these pathways depending on IDH status (Additional File 1).

### Regulation of gene expression by REST depends on *IDH1* mutation status

To evaluate an impact of REST on gene regulation in the context of *IDH1* mutation, we performed three pairwise comparisons of transcriptomic profiles between IDH-WT and IDH-MUT U87 cells. Differential analysis between IDH-WT vs IDH-MUT was performed for the untreated, siCTRL-transfected and siREST-transfected U87 IDH-WT and IDH-MUT cell lines. DEGs identified in these comparisons were intersected (Figure 3A) to validate utility of our cell line model and pinpoint silencing-specific effects. We discovered 2,943 IDH-DEGs common for the three comparisons (Figure 3A), showing a strong influence of the IDH-related phenotype on gene expression. To investigate the effect of REST knockdown on IDH-phenotype dependent genes, we focused on genes significantly altered in REST depleted cells (n=6,626) (Figure 3A, delimited in a gray circle) and compared DEGs fold changes in control and REST depleted cells. IDH-MUT/WT DEGs with similar fold changes were considered REST independent (Figure 3B). Increased DEGs (iDEGs) were defined as DEGs with a higher fold change in siREST-transfected cells (log2 FC>0.25, Figure 3C), while those with lower fold change in siREST-transfected cells (log2 FC<-0.25) were considered decreased DEGs (dDEGs). We defined DEGs as iDEGs when they had log2 FC difference between siREST and siCTRL above 0.25 and dDEGs as those that had log2 FC lower than - 0.25 (Figure 3C).

**Figure 3.**
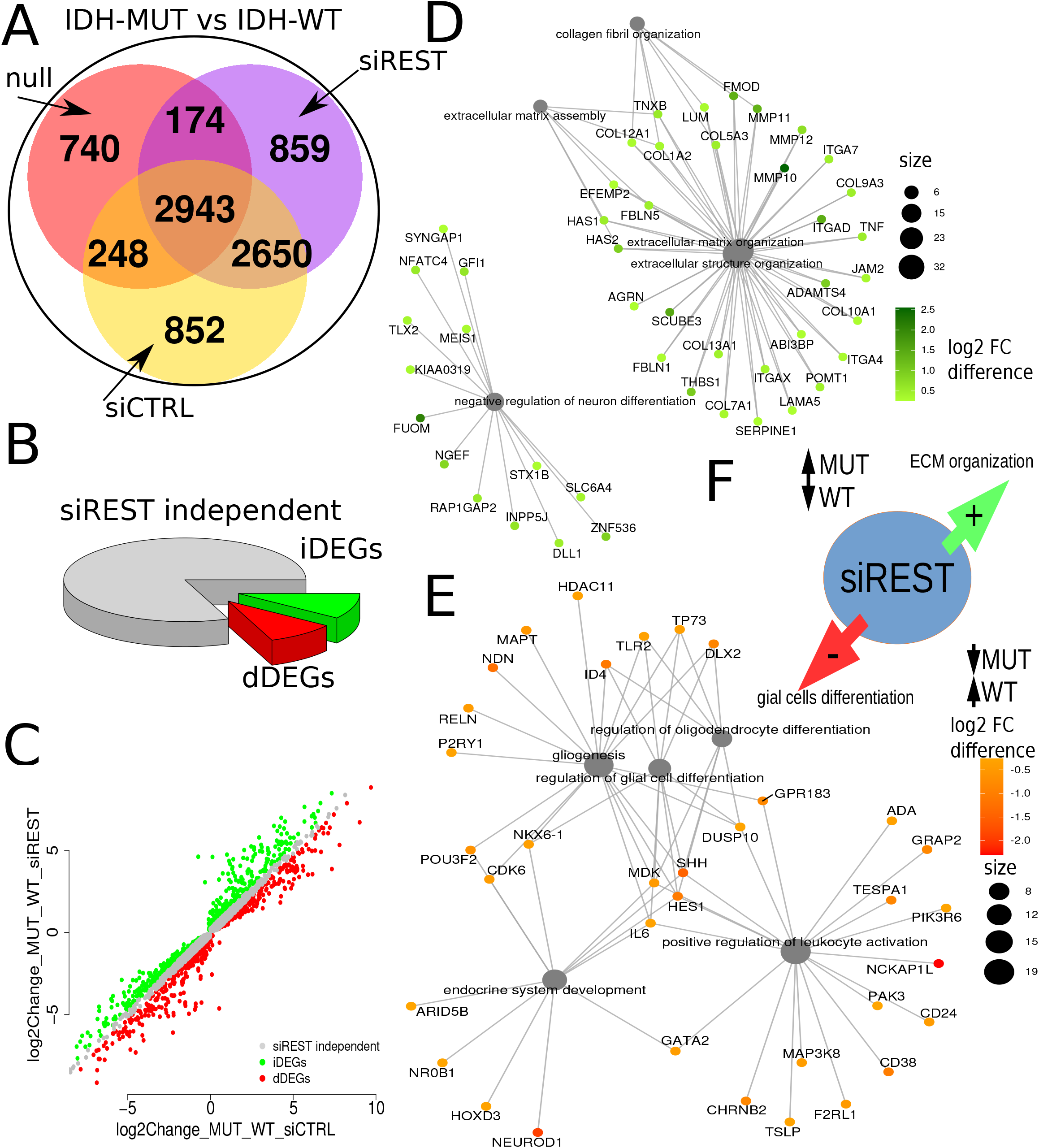
REST knockdown differentially affects expression of genes involved in extracellular matrix (ECM) organization and glial cell differentiation in IDH-WT and IDH-MUT U87 glioma cells. **(A)** Venn diagram showing the overlap of the differentially expressed genes between IDH-WT vs IDH-MUT in untreated, siREST or siCTRL-transfected U87 glioma cells. Grey circle marks DEGs in the siREST IDH-WT and IDH-MUT cells. **(B)** Subsets of IDH-DEGs altered (red, green) or unchanged in REST depleted cells (gray). **(C)** Comparison of the fold change difference in gene expression in the cells with wild type or mutated IDH, following siREST/siCTRL transfection. In siREST-transfected cells, a group of DEGs showed a shift in log2 fold change (log2 FC) between IDH-MUT and IDH-WT in siREST compared to siCTRL-transfected cells. Shift of IDH-MUT vs IDH-WT log2 FC in siREST transfected cells when compared to siCTRL transfected cells was either up (green, increased DEGs, iDEGs) or down (red, decreased DEGs, dDEGs). Difference in log2 FC was assumed significant when log2 FC shift between IDH-MUT and IDH-WT comparisons in siREST and siCTRL was >0.25 and adjusted p-value <0.05. **(D)** Gene Ontology Biological Processes (GO BP) pathways analysis of the iDEGs, showing genes associated with ECM matrix organization and negative neuronal differentiation. **(E)** Gene Ontology Biological Processes (GO BP) pathways analysis of the dDEGs, showing genes associated with glial differentiation and immune/endocrine pathways. **(F)** Graphical summary showing the opposite effect of REST knockdown in IDH-WT and IDH-MUT on the expression of genes related to ECM organization and cell differentiation.

The GO BP pathway analysis performed on these selected genes demonstrated that iDEGs showed the enrichment in genes involved in extracellular matrix (ECM) organization and negative regulation of neuron differentiation (Figure 3D). Within these pathways, the highest log2 FC increase was observed in several integrin (*ITGAD*) and metalloproteinases (*MMP10*) encoding genes. Moreover, the enrichment in genes related to reproductive, developmental, glial differentiation pathways and positive regulation of leukocyte activation was found (Figure 3E). Increased expression of genes related with ECM in IDH-MUT REST depleted cells and decrease in expression of these genes in REST depleted IDH-WT cells, contrasted with decreased expression of glial differentiation related genes in IDH-MUT REST depleted or increased expression of these genes in REST depleted IDH-WT (Figure 3F).

These findings point to a potential role of REST in a switch between ECM organization and cell differentiation in cells with the IDH-related phenotype. To corroborate these observations, the transcriptomes of G4 primary glioma cell lines were analyzed [20]. In general, differential expression between IDH-MUT (2 G4 astrocytomas) vs IDH-WT (10 G4 GBMs) was confirmed only for genes downregulated in IDH-MUT (Additional File 1A). Therefore, only down-regulated ECM-regulated genes from Fig 3D were visualized on z-score heatmap (Additional File 1B) for IDH-MUT vs IDH-WT in U87 cell lines and for primary IDH-MUT vs IDH-WT G4 gliomas (Additional File 1C). Significant differences between IDH-MUT and IDH-WT cell lines were obtained for 13 out of 18 genes (Additional File 1C), and bootstrapping confirmed significance of this result (Additional File 1D, 1.2% probability of obtaining such result by chance).

### The effect of REST knockdown on U87 glioma cell invasiveness and expression of genes associated with extracellular matrix depends on the *IDH* mutation - related phenotype

We tested whether the enrichment in Gene Ontology biological pathways related to ECM organization and cell differentiation observed in REST depleted glioma cells coexisted with changes in the invasiveness and viability of the cells. Viability of both IDH-WT and IDH-MUT U87 glioma cells was not significantly affected by REST knockdown at 12-, 24-, 48- or 72 hours post-transfection as measured with the PrestoBlue assay (Figure 4A). The invasion of REST depleted U87 IDH-WT cells quantified with Matrigel assay increased by 75% compared to control cells (Figure 4B, migrating cells: siCTRL: 908.8 ± SEM=460; siREST: 1597± SEM=622.3).). The opposite trend, observed in U87 IDH-MUT, was not statistically significant. Invasiveness of the untreated glioma cells was strongly influenced by their IDH mutation status (Figure 4B, migrating cells IDH-MUT: 3472 ± SEM=324.7; IDH-WT: 1019 ± SEM=490.6). DEGs whose fold change between U87 IDH-MUT and U87 IDH-WT was increased by REST (iDEGs, Figure 3D) included a group of genes implicated in the ECM organization pathway (Figure 4C). To validate if these genes depend on REST in the tumor transcriptome dataset, their expression was correlated with *REST* expression using TCGA glioma dataset (G2-G4). The majority of them significantly correlated with *REST* either in G4 (Additional File 3, left panel), LGG IDH-MUT (Additional File 3, middle panel) or in LGG IDH-WT (Additional File 3, right panel), supporting the notion that REST is required to regulate ECM-related gene expression, but its targets may differ depending on the IDH-MUT status. Finally, using tumor methylome data of IDH-WT and IDH-MUT (G2/G3) gliomas, hereafter “glioma Atlas” [18], we found that 16 out of the 32 identified ECM related genes had significantly differential DNA methylation in gene promoters (Figure 4D) and 11 within gene bodies (Figure 4E).

**Figure 4.**
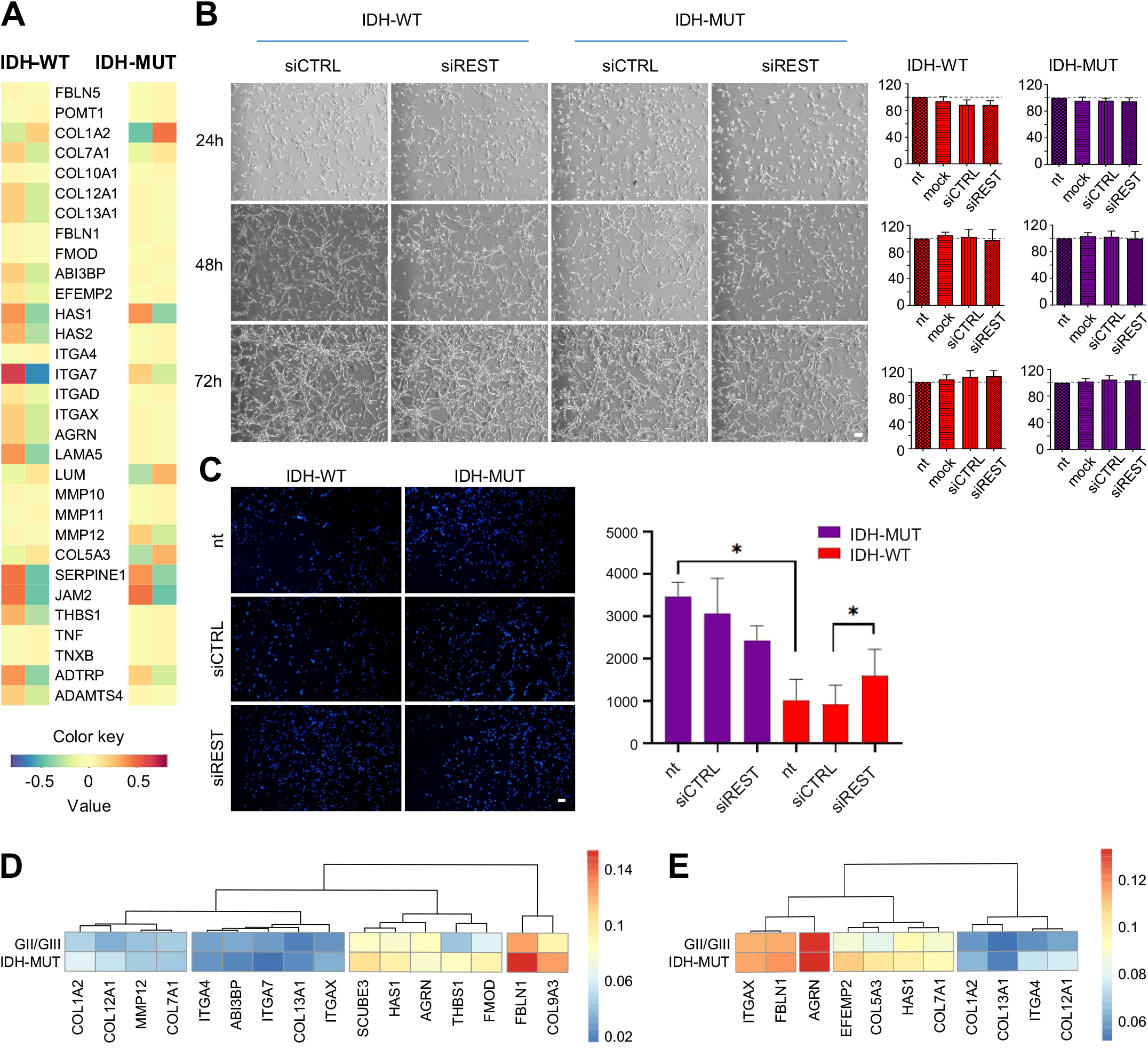
REST knockdown affects glioma cell invasiveness and expression of genes implicated in cell migration/invasion. **(A)** Bright field microscopy images of IDH-WT and IDH-MUT U87 cells 24-, 48- and 72 hours after siRNA transfection; scale bar = 200 µm. Cell viability after 24-, 48- or 72 hours of REST silencing measured with PrestoBlue assay. Data are represented as mean ± SEM, n=3, Wilcoxon matched-pairs signed rank test, two-tailed (p>0.05). Dotted line at 100% denotes a viability of mock-transfected cells. **(B)** Invasiveness of IDH-WT and IDH-MUT U87 cells measured with Matrigel invasion assay. The cells were either not treated (nt) or transfected with siCTRL or siREST. The fluorescence microscopy images (scale bar = 200 µm) show representative fields of Matrigel inserts and the bar plot shows quantification of the migrating cells. Data are presented as mean ± SEM, n=6, *p<0.05, Wilcoxon matched-pairs signed rank test. IDH-WT: siCTRL: 908.8 ± SEM=460; siREST: 1597± SEM=622.3. Not treated cells: IDH-MUT: 3472 ± SEM=324.7; IDH-WT: 1019 ± SEM=490.6; p=0.0313. **(C)** Extracellular matrix organization GO BP-related IDH-DEGs modulated by siREST (selected from functional analysis from Figure 3C) presented as a heatmap for IDH-WT (left panel) and IDH-MUT (right panel) siREST (right column in each panel) vs siCTRL (left column in each panel) U87 glioma cell lines. **(D)** Hierarchical clustering based on mean DNA methylation level within promoters (TSS - 2000/+500 bps) of ECM genes whose DNA methylation was significantly different (FDR< 0.05) between IDH-MUT and IDH-WT (G2/G3) tumor samples deposited in Glioma Atlas. **(E)** Description as in (D) but for genes with significantly differential DNA methylation in gene bodies.

### Characterization of REST ChIP-seq peaks in IDH-WT and IDH-MUT U87 cells

Chromatin immunoprecipitation followed by sequencing (ChIP-seq) was employed to identify REST binding sites in U87 glioma cells. The analysis revealed almost four thousand REST ChIP-seq peaks out of which 2,647 were common in IDH-WT and IDH-MUT cells, while 114 were specific to IDH-MUT and 1,077 to IDH-WT cells. REST-ChIP-seq peaks were annotated to genes (hereafter referred to as REST targets). Consistently, the majority of REST targets were shared between IDH-WT and IDH-MUT cells (n=1,674), but 85 genes were specific to IDH-MUT and 860 IDH-WT cells (Figure 5A). Most of the IDH-MUT specific peaks were located in the intergenic or intronic regions, while the IDH-WT were located mainly in gene promoter regions (Figure 5B). Both, the lower number of binding sites identified in IDH-MUT (Figure 5A) and more distal IDH-MUT specific binding sites localization from transcription start site (Figure 5B) indicates that the role of REST in IDH-MUT is different than in IDH-WT.

**Figure 5.**
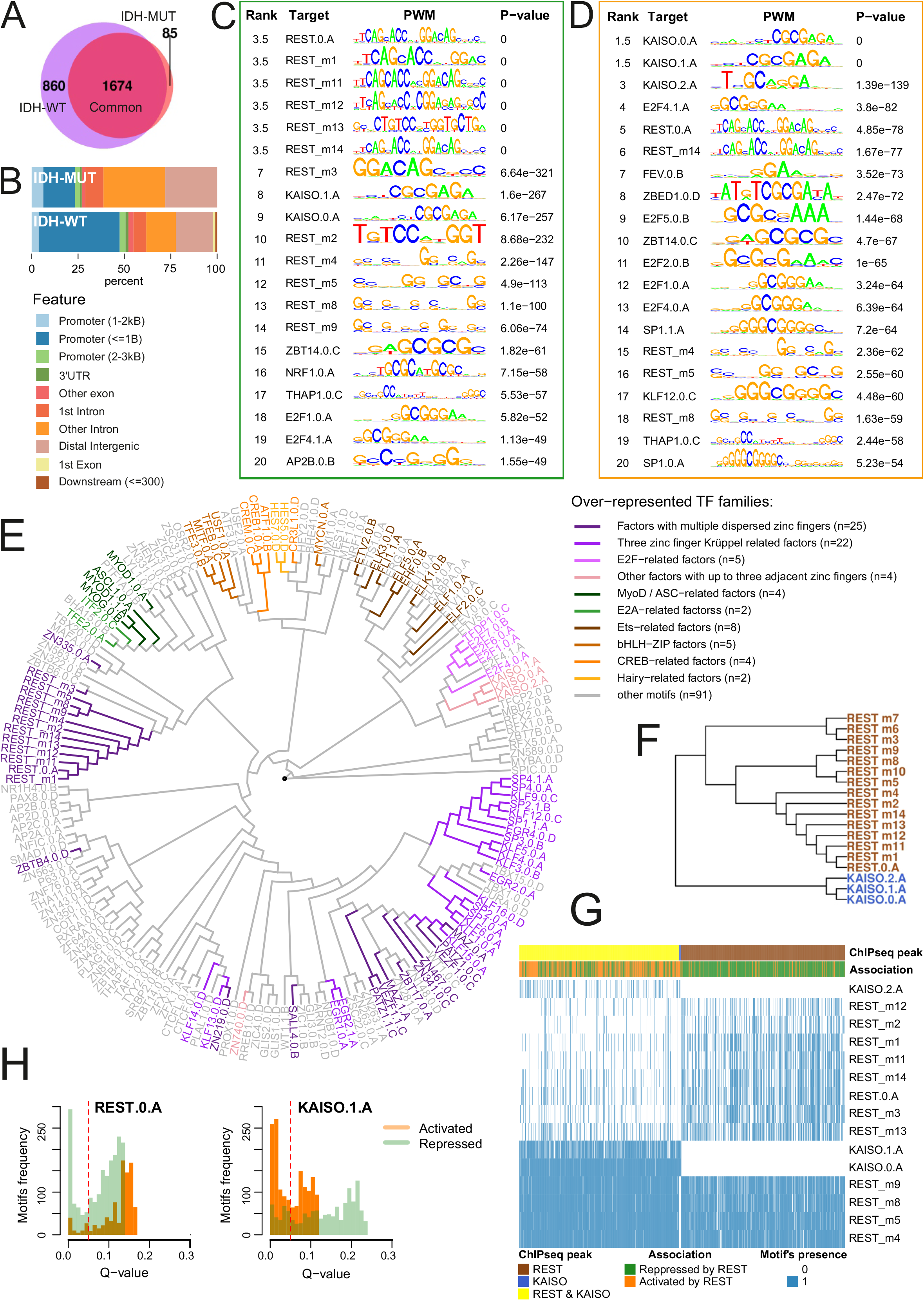
Characterization of REST-ChIP-seq peaks and their target genes. **(A)** Intersection of genes assigned to REST-ChIP-seq peaks in IDH-WT and IDH-MUT U87 cells. Peaks were assigned to genes following the R ChIPseeker library assignment to the promoter region. **(B)** Annotation of identified REST-ChIP-seq peaks to genomic regions. **(C-D)** Ranking of the TOP 20 TF motifs identified in the sequences of the REST-ChIP-seq peaks assigned to genes repressed by REST **(C)** or activated by REST **(D)**. Briefly, *REST* expression was correlated with the genes to which REST-ChIP-seq peaks were assigned (called REST targets) using TCGA dataset (data from Figure 5A). Based on the correlation results between *REST* gene expression and REST targets, the genes were divided into repressed or activated by REST. If correlation was statistically significant (adjusted p value <0.05) and correlation coefficient was positive, a gene was assigned as activated by REST, while coefficient was negative, a gene was assigned as repressed by REST. **(E)** Hierarchical tree of TF motifs for enriched TF families based on PWMs. Shades of green represent motifs from TF protein families overrepresented in REST ChIP-seq peaks unique for repressed REST targets; orange - activated; magenta - motifs overrepresented in the group of motifs present in both repressed and activated REST targets. **(F)** REST and KAISO (ZBTB33) motifs clustering based on PWMs. **(G)** Hierarchical clustering of REST peaks according to the identified KAISO and REST motifs. Color-coded bars show the association of a REST peak and its target gene, impact on gene expression (repressed or activated by REST) and the presence of REST and/or KAISO motifs. **(H)** Q-value and frequency relations for selected KAISO (ZBTB33) and REST motifs within REST-ChIP-seq peaks assigned to genes activated or repressed by REST. To highlight the pattern, bar plots show the full distribution of q-values with a red dashed line indicating significance cut-off point.

The REACTOME pathway analysis of genes annotated to REST peaks indicated the gene enrichment in ten pathways common to IDH-WT/MUT and eleven pathways specific to either IDH-WT or IDH-MUT REST targets (Additional File 4A). Ten of these pathways were specific to IDH-WT, and they were related to transcription, translation, nonsense mediated decay, voltage gated potassium channels, infectious disease, and heat shock factor 1 activation. The only pathway significantly enriched in targets specific to IDH-MUT was: “Interaction between L1 and ankyrins” (Additional File 4A).

Next, we examined sequences of REST-ChIP-seq peaks assigned to genes specific to IDH-WT (n=860), IDH-MUT (n=85), or common to both cell types (n=1,674) (Figure 5A) to find transcription factor (TF) binding sites within these regions and identify TFs whose binding might depend on the IDH-related phenotype. Using the EnrichR tool and the ENCODE ChIP-seq dataset, we identified TFs overlapping within REST-ChIP-seq peaks (Additional File 4C). In particular, we found that ENCODE ChIP-seq peaks of KAISO known also as ZBTB33 (Zinc Finger and BTB Domain Containing 33) had the strongest enrichment within the REST-ChIP-seq peaks specific for IDH-WT cells (Additional File 4C, middle panel). In IDH-MUT specific REST-ChIP-seq peaks, only REST motifs were identified (Additional File 4C, middle panel), while in the REST-ChIP-seq peaks common for the IDH-WT and IDH-MUT, REST motifs appeared at the top positions of the enrichment ranking, but at the lower positions, KAISO motifs were present as well (Additional File 4C, top panel).

KAISO transcription factor has been reported in several different human cancers functioning as a tumor suppressor or oncogene [31]. KAISO function seems to be highly context-dependent [32]. It binds to methylated CpGs in two motifs containing a consensus sequence 5’-CGCG-3’ (Figure 5C and D: KAISO.0.A and KAISO.1.A) and to unmethylated C in the motifs with another consensus sequence 5’-CTGCNA-3’ (Figure 5C and D: KAISO.2.A) [31].

To identify specific TF motifs, present within the REST ChIP-seq peaks specific for IDH-WT or IDH-MUT, or common for both IDH-WT and IDH-MUT, we investigated the peak sequences using all available position weighted matrices (PWMs) deposited in the **HO**mo sapiens **CO**mprehensive **MO**del **CO**llection (HOCOMOCO) database (version 11) and additional 14 REST PWMs from the ENCODE dataset. Using the R PWMEnrich package and the FIMO tool [30], we identified 14 motifs in IDH-MUT specific REST-ChIP-seq peaks, 70 in IDH-WT-specific and 120 in the common peaks (Additional File 4B). Ten motifs were shared among the common, IDH-WT and IDH-MUT REST ChIP-seq peaks (Additional File 4B). Focusing on KAISO motifs, we discovered that all the three HOCOMOCO motif variants (0.A, 1.A and 2.A) were present within the common and the IDH-WT-specific REST-ChIP-seq peaks, but not in the IDH-MUT-specific peaks (Additional File 4D).

We hypothesized that changes in DNA methylation resulting from an IDH mutation may affect binding of REST to its respective sites. Therefore, we examined DNA methylation levels within all the REST-ChIP-seq peaks detected in the U87 cells. We analyzed methylome data from the glioma Atlas [18] using the DiffMeth tool. DNA methylation level was determined in G2/G3 IDH-MUT, G2/G3 IDH-WT and G4 IDH-WT glioma samples. Pairwise comparisons of DNA methylation between different glioma samples were performed separately for the sequences in U87 REST-ChIP-seq peak assigned as common, IDH-WT or IDH-MUT specific (Additional File 5A-C). A similar percentage of common, IDH-WT and IDH-MUT specific peaks was differentially methylated: 8%, 6.9%, 5.5% respectively. The largest number of differentially methylated peaks was detected between a pair of G2/G3 IDH-MUT vs G4 IDH-WT samples (Additional File 5A-C, middle column). Next, to identify the effect of IDH-related phenotype, we focused on peaks differentially methylated between IDH-WT and IDH-MUT G2/G3/G4 samples (Additional File 5A-C, left column). In majority of cases IDH-MUT samples had higher DNA methylation (Additional File 5D-F), showing that rather IDH mutation status then peak origin impacts more on differential DNA methylation pattern of REST-ChIP-seq peaks in glioma tumors.

### Variability of TF motifs within the REST ChIP-seq peaks

Most REST-repressed genes were enriched in the GO biological pathways related to neuronal functions, confirming its canonical role as a repressor of neuronal genes in non-neuronal cells (Additional File 6A, left panel). On the other hand, pathways related to the REST-activated targets were more diverse (Additional File 6A, right panel).

To determine the exact locations of the detected TF motifs in REST-activated and REST-repressed genes, we used the FIMO tool and identified 145 motifs for 119 TFs within the REST peaks assigned to REST-repressed genes and 140 motifs for 115 TFs within the peaks assigned to REST-activated genes (q-value ≤0.05). Hierarchical clustering tree of these TF motif PWMs revealed grouping of REST-repressed and REST-activated gene specific motifs (Additional File 6B). The same tree, exhibiting TF motifs sequence similarities, was used to visualize protein families of TFs detected within REST peaks paired with REST-activated or repressed genes (Figure 5E). The motifs identified in both REST-repressed and activated targets included E2F transcription factor family motifs, which is a classic TF involved in glioma progression [33,34].

Among the motifs enriched in REST-ChIP-seq peaks of REST-repressed genes, were targets for ASCL1, MyoD and E2A-related factors that are engaged in cell differentiation and proliferation, including neuronal (ASCL1) and oligodendrocyte (E2A) differentiation [35], mesenchymal cell proliferation (MyoD) and growth inhibition [36]. In cancer, these TFs act as activators (ASCL1) or inhibitors of the cell cycle progression (E2A) [37]. Motifs overrepresented in REST-activated genes were targets for CREB, Ets family proteins, bHLH-ZIP, and Hairy-related factors. Among others, CREB regulates transcription of the genes coding for a proto-oncogene c-Fos [38] and neuropeptide VGF [39]. The latter was one of the stronger upregulated genes upon REST knockdown (Figure 2D). Ets family proteins are activated by Ras-MAP kinase signaling pathway and have been implicated in tissue differentiation and cancer progression [40]. Hairy-related proteins typically function as DNA-binding transcriptional repressors that have been shown to inhibit Notch activated *a-actin* [41] and control differentiation [42]. Overrepresentation of the motifs for these TFs in the vicinity of REST binding sites in REST-repressed genes shows a potential contribution of their pathways to the diverse REST effects and its impact on tumorigenesis and patient survival.

To identify the most prevalent TF motifs within the REST peaks associated with REST-activated or REST-repressed genes, we performed TF motif scans using PWMs. The top of the ranking of the highest scoring motifs for the REST-repressed genes contained mainly REST motifs, as expected (Figure 5C). However, in the ranking of TF motifs in REST-activated genes, the first three positions were occupied by KAISO (Figure 5D). TF binding motif sequence logos generated for both KAISO and REST PWMs (Additional File 6F, G) and hierarchical clustering of all discovered PWM sequences (Figure 5E) as well as REST and KAISO PWMs alone (Figure 5F) showed dissimilarity between these motifs, hence their co-occurrence within the REST ChIP-seq peaks was not due to the PWMs similarity. However, based on the distribution of the statistical significance of the occurrence of TF motifs, we may assume that KAISO motifs are more frequent in the promoters of REST-activated than REST-repressed genes, whereas REST motifs display a somewhat reverse pattern (Figure 5H). The results of CentriMo analysis showing the probability of REST or KAISO motif presence in the gene promoters corroborate this observation, as we found more frequent occurrence of KAISO motifs in genes activated by REST (Additional File 6C) than in the genes repressed by REST (Additional File 6D). For the REST motifs in the REST-repressed genes the distribution was similar to KAISO motifs (Additional File 6D-E) while for the REST-activated genes the probability was below statistical significance.

To determine a degree of REST-ChIP-seq peaks similarity according to the detected REST and KAISO motifs, each peak was represented by a binary vector where a given REST or KAISO motif was present (value=1) or absent (value=0), and a hierarchical clustering on these vectors was performed (Figure 5G). In general, two clusters of peaks were discovered: 1) represented mostly by REST and KAISO motifs and associated with gene activation or 2) represented only by REST motifs and mostly associated with gene repression (Figure 5G). Among the reported REST peaks (n=1,523), 63% were related to repressed targets and 37% to activated targets. In the majority of peaks related to REST-repressed genes (69.5%), only REST motifs were found. In contrast, in the majority of peaks related to activated genes (81%) both REST and KAISO motifs were present. Our results imply that KAISO motif occurrence or binding may be an important factor in REST-guided gene activation, but not in gene repression.

### Distinct DNA methylation within REST and KAISO motifs of activated and repressed REST targets

DNA methylation contributes to activation or repression of transcription. DNA methylation can influence TFs binding and activity when occurring in the vicinity of their binding sites. While in the previous analysis (Additional File 5) we verified DNA methylation of the whole 200 bp sequences of REST-ChIP-seq peaks, here we focused on DNA methylation levels of REST and KAISO motifs detected within the REST-ChIP-seq peaks assigned to REST-repressed or activated targets to verify whether DNA methylation may play a modulatory role in REST-guided gene regulation. We studied methylated cytosines in the CpG context with a minimal coverage of 10 reads in the G2/G3 IDH-MUT, G2/G3 IDH-WT and G4 IDH-WT glioma methylomes deposited in glioma Atlas [18]. These methylomes, covering millions of sites at 1 bp resolution, were intersected with coordinates of KAISO and REST motifs derived from U87 REST-ChIP-seq. In result, we obtained 23,614 unique cytosines within TF motif sequences. The majority of cytosines occurred within either REST or KAISO motifs, but 2,588 were located within overlapping sites of REST and KAISO motifs. When a cytosine was located in multiple motifs of a single TF (due to motifs overlapping), its beta value (β) was only counted once. When a cytosine was located in overlapping motifs of different TFs (REST or KAISO), its beta value was counted separately for each TF motif. For each cytosine position, a mean methylation (β-value) across samples was computed and discretized into low [0-0.2], medium (0.2-0.6] or high (0.6-1]. Low β-values correspond to DNA hypomethylation, whereas high β-values indicate DNA hypermethylation. REST motifs in the REST-ChIP-seq peaks lacking KAISO motifs and assigned to the repressed targets, were significantly enriched (p <2.2x10^-16^) in medium- and hypermethylated cytosines (Figure 6A, Additional File 7). In these REST motifs, the percentage of hypermethylated cytosines was around three times higher than within REST-ChIP-seq peaks assigned to the activated REST targets (Figure 6A-B, Additional File 7).

**Figure 6.**
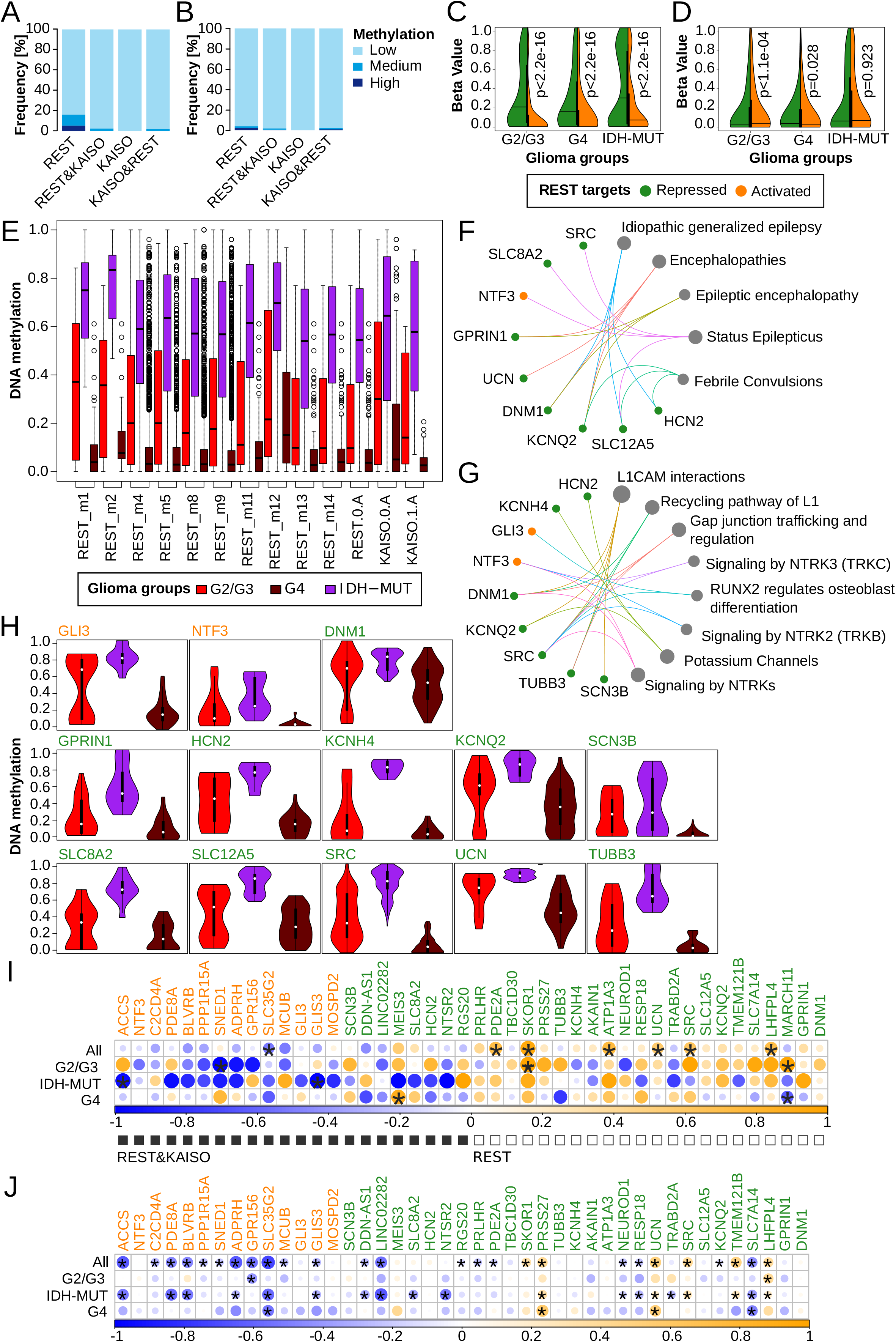
DNA methylation distribution at the selected REST-ChIP-seq peaks and its influence on REST targets. **(A)** DNA methylation of REST or KAISO motifs within REST-ChIP-seq peaks assigned to REST-repressed targets using IDH-MUT (G2/G3) and IDH-WT (G2/G3/G4) tumor samples from glioma Atlas. Description represents: REST – methylation of REST motifs of peaks lacking KAISO; REST&KAISO – methylation of REST motifs of peaks containing both REST and KAISO motifs; KAISO – methylation of KAISO motifs present in REST ChIP-seq peaks with identified KAISO but not REST motif; KAISO&REST – methylation of KAISO motifs of peaks containing both REST and KAISO motifs. **(B)** DNA methylation of REST or KAISO motifs within the REST-ChIP-seq peaks assigned to REST-activated targets based on IDH-MUT (G2/G3) and IDH-WT (G2/G3/G4) tumor samples from glioma Atlas. Description as in (A). **(C)** Distribution of REST motif DNA methylation in the REST ChIP-seq peaks containing only REST motifs and assigned to the repressed or activated REST targets based on glioma Atlas. **(D)** Distribution of REST motif DNA methylation in the REST ChIP-seq peaks containing both REST and KAISO motifs and assigned to the repressed or activated targets based on glioma Atlas. **(E)** Cumulative distribution of DNA methylation in individual motifs. There are 601 sites containing REST or KAISO motifs with significantly different methylation among gliomas from glioma Atlas. **(F)** Human disease pathways enriched among 47 REST targets. REST-ChIP-seq peaks assigned to these targets had at least one differentially methylated REST or KAISO motif. **(G)** Description as in (F) but for REACTOME pathways. **(H)** Distribution of DNA methylation (glioma Atlas) of REST or KAISO motifs within REST-ChIP-seq peaks assigned to the genes present in enriched pathways shown in (F) and (G). **(I)** Correlation between expression of a REST-target gene and mean DNA methylation of a motif assigned to its promoter. Correlations were performed using the glioma Atlas dataset. Black squares mark REST-targets having REST and KAISO motifs in the REST ChIP-seq peaks, while white squares mark those in which only REST motifs were detected. **(J)** Correlation between expression of a REST-target gene and mean DNA methylation of its promoter. Analyses performed using the data from TCGA dataset.

Since a vast majority of cytosines were hypomethylated (Figure 6A-B), methylation variance across G2/G3 IDH-MUT, G2/G3 IDH-WT and G4 IDH-WT gliomas was computed for each cytosine within REST and KAISO motifs. The obtained variance ranged from 0 to 0.05 among sites, with an average β-value of 0.0067. To explore the methylation pattern within REST motifs, we selected cytosines with a higher variance than the average. We found that DNA methylation of REST motifs located within REST-ChIP-seq peaks lacking KAISO motifs was significantly higher in peaks assigned to REST-repressed targets than in peaks assigned to REST-activated targets across all glioma groups (Figure 6C). DNA methylation within REST motifs of REST-ChIP-seq peaks containing both KAISO and REST motifs showed significantly higher DNA methylation in peaks assigned to repressed REST targets only in G4 gliomas (Figure 6D). In G2/G3 samples the methylation pattern was the opposite and in IDH-MUT no significant differences were detected (Figure 6D).

The number of sites with predicted REST and/or KAISO motifs within a single REST-peak was high, which did not allow for unequivocal determination which of predicted motifs was actually used for TF binding. Thus, we studied DNA methylation patterns within individual motifs for the REST or KAISO TFs across gliomas using the DiffMeth tool and methylomes of G2/G3 IDH-WT, G2/G3 IDH-MUT and G4 IDH-WT gliomas from the glioma Atlas [18]. Initially, over 70,000 REST and KAISO motif sites were found within REST-ChIP-seq peaks of glioma cells (Additional File 8), and 601 of them were differentially methylated among gliomas, which accounts for 0.9% of the all identified REST and KAISO motif sites (Figure 6E). The proportion between all predicted sites of individual motifs compared to a number of differentially methylated sites was significantly different (p-value <0.01, Additional File 8). We identified six of the twelve REST motifs as having differential DNA methylation in gliomas (Additional File 8) and a majority of differentially methylated sites of REST motifs were found within REST-ChIP-seq peaks corresponding to REST-repressed targets (X-squared = 97.069, df = 13, p-value =6.119e-15).

Differentially methylated sites (n=601) among gliomas had the highest median β-value in IDH-MUT and the lowest in G4 gliomas (Figure 6E). Frequently, more than one motif showed significantly differential methylation in a single REST-ChIP-seq peak. We found that sites differentially methylated in gliomas appeared within 47 REST-ChIP-seq peaks, out of which 32 were associated with REST-repressed targets, while 15 with REST-activated targets. A complete list of REST targets paired with the REST-ChIP-seq peaks containing differentially methylated REST or KAISO motif sites is presented in Additional File 9. For these genes, we searched for associations with diseases using the DisGeNET platform and enrichment of specific biological pathways using the REACTOME database. Nine out of 47 genes were linked to disorders, including epilepsy, encephalopathies, and febrile convulsions (Figure 6F). Nine genes were associated with significantly enriched REACTOME pathways (Figure 6G). The largest number of genes (n=5), all under REST repression, were linked with L1 cell adhesion molecule (L1CAM) interactions. Several genes (n=6), all assigned as REST-repressed targets, were linked to another three pathways: recycling pathway of L1, gap junction trafficking and regulation, and potassium channels (Figure 6G). We found that the biological functions associated with L1CAM interactions and cell differentiation (*RUNX2* regulates osteoblast differentiation) are similar to pathways detected to be altered in IDH-WT and IDH-MUT REST depleted cells (Figure 3). Some of the genes defined as REST-activated targets (namely, *NTF3* and *GLI3*) and the genes containing KAISO motifs within the REST-ChIP-seq peaks (namely *GLI3, NTF3, HCN2, SCN3B* and *SLC8A2*) had a methylation pattern similar to the repressed REST targets (Figure 6H).

To verify whether differentially methylated motif sites affect expression of REST targets, we calculated correlation between mean methylation and gene expression using the glioma Atlas (Figure 6I) and TCGA datasets (Figure 6J). The methylome data in the glioma Atlas are dense, thus we were able to calculate TF motif site mean methylation and correlate it with REST target expression. Correlations were also performed on TCGA data to validate the results obtained on the glioma Atlas, but due to the low number of cytosines covered, mean methylation was computed for the entire promoters. Expression of most REST activated targets negatively correlated with site/promoter methylation levels. Interestingly, for the repressed genes, the positive correlation was more frequently detected (Figure 6I-J). The correlation results obtained on the glioma Atlas data demonstrate that a negative correlation coefficient corresponds well with KAISO motif presence in a ChIP-seq peak assigned to a gene (Figure 6I black squares) and a negative correlation with its absence (Figure 6I white squares).

### REST-ChIP-seq peaks DNA methylation of siCTRL and siREST treated IDH-WT and IDH-MUT samples of U87 cell line

REST knockdown had a modest effect on DNA methylation in the U87 cell line of a limited number of cytosines within REST ChIP-seq peaks. Out of 308816 cytosines tested in total, the difference in single cytosine methylation beta values in IDH-WT siREST and siCTRL samples was non-existent for over 99% of loci. For 79 loci the difference was below zero (lower beta values in siREST) and for 96 was above zero (higher beta values in siREST). The highest difference of beta-value was 0.08 (Additional File 10A). Similarly, for IDH-MUT, out of 317501 loci tested, the difference in DNA methylation beta values between siREST and siCTRL was identified in 54 loci as negative (lower in siREST) and in 81 loci as greater than zero (higher in siREST). Here, the highest difference of beta value of a single cytosine locus between siREST and siCTRL was 0.09 (Additional File 10B). The pattern of a difference in methylation of single cytosines between IDH-WT and IDH-MUT was fairly constant regardless of REST knockdown (Additional File 10C - comparison of WT and MUT transfected with siCTRL; Additional File 10D comparison of WT and MUT transfected with siREST). Out of all cytosines (n=833) categorized as highly methylated in IDH-MUT, only one did not match between siREST and siCTRL samples, while in cytosines having lower methylation in IDH-MUT (n=746) only two did not match between siREST and siCTRL. The observed difference in single cytosine methylation between IDH-MUT and IDH-WT was almost symmetrical for the siCTRL and siREST transfected samples (Additional File 10E), suggesting a similar effect of REST knockdown in U87-MG regardless of their IDH mutation status. Interestingly, the distribution of differentially methylated cytosines was significantly different (Chi2 test p<0.001) in comparison to the distribution of all cytosines present within REST-ChIPseq peaks (Background) across the three peak types. Similarly, the distribution of cytosines with higher or lower methylation in IDH-MUT samples differed significantly from the background showing enrichment in common and MUT-specific REST-peaks and depletion in WT-specific REST-ChIPseq peaks (Additional File 10F). It is worth mentioning that proportionally much higher enrichment of differentially methylated cytosines was found in MUT-specific REST-ChIPseq peaks (Additional File 10G).

### REST gene targets affected by REST knockdown belong to cell migration and differentiation pathways

REST knockdown may directly affect genes that are activated or repressed by REST binding (hereafter primary REST targets). REST knockdown may also have indirect effects related to altered interactions between REST and other proteins and/or downstream regulatory cascades released by primary REST targets (hereafter REST secondary targets). To separate primary from the secondary REST targets, the genes significantly affected by REST knockdown were intersected with genes assigned to REST ChIP-seq peaks. GO Biological Pathways enriched among the genes upregulated in REST depleted cells and having REST binding at their promoters’ comprised pathways related to neuronal transmission (signal release, calcium regulated exocytosis), and glial cell migration (Figure 7A, upper panel). The pathways related to the genes downregulated in REST depleted cells and whose promoters showed REST binding, included NAD biosynthetic process, regulation of mRNA splicing and hematopoietic progenitor cell differentiation (Additional File 11B).

**Figure 7.**
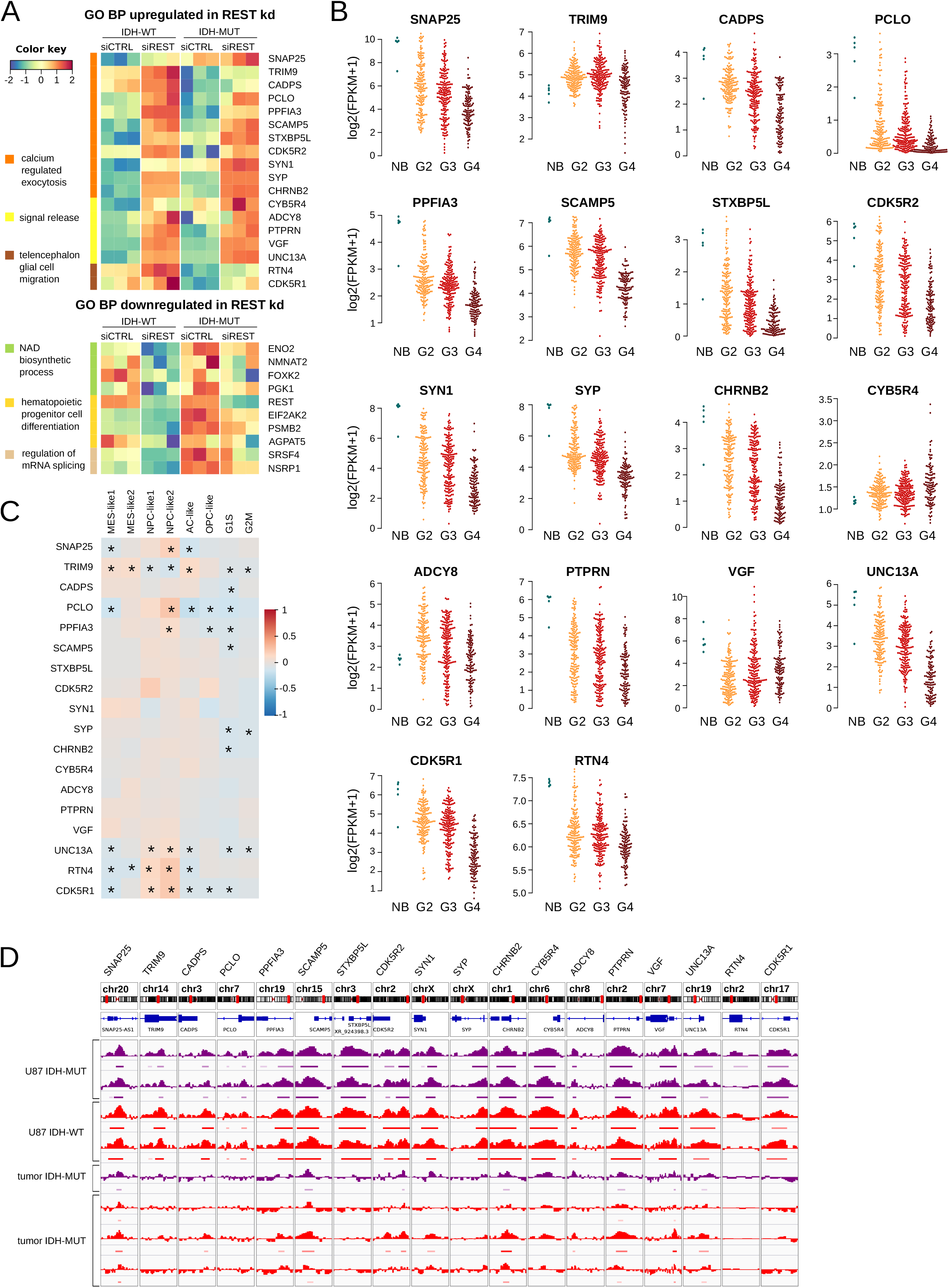
Genes affected by REST knockdown and having REST-ChIP-seq peaks belong to migration and differentiation pathways. **(A)** Upper panel: top GO BP pathways upregulated in REST depleted cells and defined as REST targets in ChIP-seq experiments were related to calcium release exocytosis, signal release and encephalon glial migration pathways. Lower panel: Main GO BP pathways downregulated in REST depleted cells and defined as REST targets in ChIP-seq experiments were related to: NAD biosynthetic process, hematopoietic progenitor cell differentiation and regulation of mRNA splicing pathways. **(B)** Expression of genes upregulated in REST depleted cells (from panel A) in TCGA LGG/GBM datasets and presented as bee swarm plots for NB (normal brain), glioma WHO G2, G3 and G4. **(C)** Expression of genes upregulated in REST depleted cells was correlated in single cell RNAseq data with cellular states defined by Neftel et al 2019. Pearson correlation was calculated, and a color key scale was used to present its values in range of -1 to 1, correlation significance was marked (*) when adjusted p <0.05. **(D)** Genomic view from Integrative Genome Viewer on genes upregulated in REST depleted cells and gliomas for each sample a histogram of reads was normalized to the input reads (bigwigg file) and bed file from ChIP-seq experiment was shown. REST ChIP-seq data on four U87 glioma cell repetitions (2x IDH-MUT and 2x IDH-WT) and 4 tumor samples (1x IDH-MUT and 3x IDH-WT) are shown. IDH-MUT samples are color-coded in purple and IDH-WT in red.

Further, a correlation of REST and its targets expression was calculated in the TCGA dataset. A majority of the genes enriched in the signal release, calcium regulated exocytosis and telencephalon glial cell migration biological pathways in REST depleted cells (Figure 7A) was similarly regulated in G4 gliomas (Additional File 11A). Nearly half of the downregulated genes in REST depleted cells related to NAD biosynthetic process, regulation of mRNA splicing and hematopoietic progenitor cell differentiation (Figure 7A), showed similar pattern in G4 gliomas (Additional File 11B). Most of the genes upregulated in the cells with REST knockdown were consistently downregulated in gliomas compared to normal brain samples, with expression decreasing in the direction from WHO G2 and G3 to G4 (Figure 7B). Genes downregulated in REST depleted cells did not show this pattern in TCGA gliomas (not shown). Correlation of the REST up- or down-regulated genes with glioma cellular states described by Neftel et al. 2019 showed that many of the genes correlated with cellular states that recapitulate neural progenitor-like (NPC-like), oligodendrocyte progenitor-like (OPC-like), astrocyte-like (AC-like) or mesenchymal-like (MES-like) states and/or states characteristic for cycling cells related to G1S or G2M cell cycle transition phases (Figure 7C). Many genes upregulated in REST depleted cells show significant positive correlation with NPC-like states and a negative correlation with G1S and G2M states, suggesting rather non-cycling cells (Figure 7C). Finally, the genes upregulated in REST depleted U87 glioma cells show a concordance of their REST binding sites with these detected directly from glioma tumors, collected within the study (Figure 7D), which was not the case among genes downregulated in REST depleted cells (not shown).

## Discussion

We performed a comprehensive study identifying transcriptional targets of REST transcription factor and complete REST regulatory networks in U87 glioma cells with a different IDH status. These findings were validated in glioma patient samples of different malignancy and public TCGA datasets. Using a large spectrum of computational methods, ChIP-seq, RNA-seq, DNA methylation and RNAi mediated REST knockdown in U87 cells with a wild type or mutant *IDH1*, we addressed a complex role of REST in gliomagenesis. Recognition of the importance of IDH1/2 mutations in progression of diffuse gliomas advanced our understanding of glioma biology, however the full impact of a state of DNA and histone hypermethylation, leading to the CpG island methylator phenotype, on gene regulatory networks and cell functions is less clear. Our study demonstrates that REST-regulated gene networks in gliomas are dependent on the *IDH* mutation status, which determines a selection of REST dependent genes involved in ECM organization, glioma invasion and cell differentiation. We uncovered a putative cooperation between REST and KAISO in determining REST target repression or activation. Our results point to REST as a valid target in anti-glioma therapy.

Exploration of TCGA datasets showed increased *REST* expression in malignant gliomas and in IDH-mutant gliomas. High *REST* expression has been reported as a negative prognostic factor for survival in GBM [7] and medulloblastoma patients [43]. In mice, expression of REST in glioma stem cells (GSCs) was negatively correlated with survival and considered as a critical factor in maintenance of their self-renewal [2]. While in a joint cohort of lower grade gliomas and GBM *REST* expression was inversely correlated with patients survival as previously reported [2,15,44], we found that its high expression is an unfavorable prognostic factor in LGG with the *IDH* mutation, but a favorable factor in GBM (Figure 1). Finding a positive correlation of *REST* expression with survival of GBM patients appears surprising and requires more studies.

Several studies reported that REST acts as an oncogene in gliomas, promoting cell proliferation and invasion [2,15]. REST expression was associated with high tumor aggressiveness and invasiveness, as well as chemotherapy resistance [2,15,44]. However, the underlying gene regulatory networks have not been elucidated yet. REST knockdown in U87 glioma cells affected many biological pathways. Numerous genes linked to neuronal functioning were upregulated, while genes linked to cell proliferation were downregulated in REST depleted cells, regardless of the *IDH* status (Figure 2F). However, a large subset of DEGs between IDH-WT and IDH-MUT U87 glioma cells were differently affected by REST knockdown (Figure 3A). We found those genes belonged to GO biological processes related to ECM organization and glial/neuronal cell differentiation. Contrary to Zhang et al observations [16], REST knockdown did not affect cell viability in our model (perhaps due to experimental setup differences), but it did influence cell invasiveness in IDH phenotype-specific manner. We also observed a much higher invasion of the IDH-MUT compared to IDH– WT cells. These observations raised a question on how the changes in DNA methylation resulting from the *IDH* mutation affect REST targets and processes in which they are involved.

Using U87 IDH-WT and IDH-MUT isogenic cell lines and tumor samples DNA methylation data allowed us to address the issue of how DNA hypermethylation affects sets of REST regulated genes depending on IDH mutation status. REST-ChIP-seq data on human U87 glioma cells integrated with the results of transcriptomic changes in REST depleted U87 cells allowed a precise definition of direct and indirect REST targets. Scrutinizing REST-ChIP-seq peaks we found REST binding motifs co-occurring with the binding sites for other TFs, among which KAISO was identified as an important partner in gene regulation. Interestingly, depending on the co-occurrence of REST and KAISO binding sites the effects on transcription varied and different GO biological pathways were found to be regulated. We confirmed that genes selected as REST targets in cultured glioma cells, were co-expressed in a REST dependent manner in TCGA glioma datasets.

The identified REST gene regulatory networks and biological functions agree with observations of a key role of REST as a repressor of neuronal genes in non-neuronal cells. REST depletion promotes neuronal differentiation [45], while REST stabilization promotes maintenance of NPCs [46]. However, ablation of REST expression to 1% of wild type levels appeared to impede the development of NSCs, NPCs and neurons [47]. REST regulates the timing of neural progenitor differentiation during neocortex development [48]. REST and CoREST modulate not only neuronal but also glial lineage elaboration [49]. Both REST and CoREST were found to target genes encoding factors involved in mediating glial cell identity and function [50]. Blocking the function of REST suppressed the nitric oxide-induced neuronal to glial switch in neural progenitor cells [51]. REST expression was upregulated and sustained by BMP signal activation in the course of astrocytic differentiation of NPCs, which restricted neuronal differentiation [52]. REST function is also required for the differentiation of OPCs into oligodendrocytes [53].

REST-depleted neural stem cells are defective in adherence, migration and survival [47]. Yet, REST involvement in cell migration and invasion can be seen as ambiguous. On one hand, REST blocks NPC radial migration during neurogenesis [48] and acts as a cell migration repressor in microglia [54]. However, medulloblastoma cells overexpressing REST migrated faster in wound-healing assay compared to controls [55]. One possible explanation of this seeming discrepancy is that cell migration in the brain can be of different types: tangential or radial [56] and REST may contribute differently to each of these types. In addition, the migratory pathway affected may be reflective of the differentiation status of the tumor. Downregulation of REST in glioblastoma cells inhibited cell migration and proliferation [16]. In our dataset we did not observe change in cell proliferation after REST knockdown with siRNA, perhaps due to a difference in experimental procedures between Zhang’s et al [16] and us (transfection procedure, siRNA used, type of proliferation assay applied). However, we observed that IDH-MUT glioma cells are considerably more invasive (Figure 4B), which is consistent with data on IDH-MUT tumors [57–59]. In this study, REST knockdown increased expression of ECM related genes (Figure 4C) and invasion of IDH-MUT glioma cells (Figure 4B) This coincided with changes in promoter and gene body methylation in more than half of the cases (Figure 4D, E). We propose that REST-regulated genes from the ECM related biological pathway could contribute to the increased invasiveness of IDH-MUT gliomas.

The analysis of the REST-binding sites in ChIP-seq peaks shed light on different REST activity depending on the IDH status. The differences in the number of ChIP-seq peaks (Figure 5A) and in genomic distribution of DNA binding sites (Figure 5B) between IDH-WT ad IDH-MUT suggest a stronger transcriptional regulation by REST (including both repression and activation) in IDH-WT glioma cells. Some genomic locations of REST ChIP-seq peaks were exclusive to IDH-WT or IDH-MUT cells, further adding up to the possible differences in REST regulation in IDH-WT and IDH-MUT. Moreover, the REST ChIP-seq peaks in IDH-WT and IDH-MUT cells contained differing sets of other transcription factor binding motifs (Additional File 4). Different compositions of other TF motifs were also detected between peaks assigned to REST-repressed and -activated targets (Figure 5 and S6) suggesting differences in gene regulation by specific factors [60–62]. Increased *REST* expression in G4 gliomas and decreased in IDH mutant gliomas may result in REST binding to the sites that are otherwise occupied by other TFs with a different mode of action. Potential competitors of REST in regulation of its targets in IDH-WT and IDH-MUT cells included KAISO (ZBTB33), a methylation-sensitive TF [63,64].

KAISO is a transcriptional repressor acting as either a tumor suppressor or oncogene in various human cancers [32]. We found that KAISO TF motifs discriminate between the REST binding sites specific for IDH-WT and IDH-MUT cells. All three KAISO motifs were found in a number of REST ChIP-seq peaks specific for IDH-WT and common for both IDH-MUT and IDH-WT cells. Contrary, not a single KAISO motif was detected among the motifs present in the REST ChIP-seq peaks specific for IDH-MUT cells (Additional File 4). Depending on the co-occurrence of REST and KAISO binding sites, the effects on transcription and DNA methylation patterns varied (Figures 6A-E, 6G, 7E, 7F) and different GO biological pathways were found to be regulated (Additional File 6A,B). In addition, the KAISO-binding motif 2.A, which is bound by KAISO when unmethylated, was among the top motifs found in REST-activated but not in REST-repressed genes (Figure 5C, G). While the repressive role of REST has been well documented, its role in gene activation is less described and our findings contribute to understanding of this topic. REST gene activation in glioma cells is consistent with previous findings indicating that REST splice isoform REST4 activates gene expression in neural cells [65]. Possibly, glioma cells may hijack this neuronal cells’ specific regulation for their own purposes. One possible mechanism that has to be further elucidated is related to DNA methylation, which has an atypical relationship with targeted gene expression. Our findings suggest that REST activated genes promoters’ methylation may be positively correlated with their expression, in contrast to a common view (Figure 6 I,J). An example of enhancing REST binding by DNA methylation was demonstrated in developing mouse hearts, but that case was connected to gene repression [14].

Regulation of gene expression by REST is complex and includes a variety of cofactors, including mammalian SIN3 transcription regulator family member A (mSin3a) and REST corepressor 1 (CoREST). This complex mediates the recruitment of additional chromatin modulators, such as histone deacetylases (HDAC1/2), histone demethylases (G9a, SUV39H1, LSD1), MeCP2, and the DNA methyltransferase DNMT1, and generally leads to repression of gene expression [66,67].

In mouse ES cells, the absence of REST led to an increased DNA methylation at specific REST binding sites, suggesting that REST binding may lead to hypomethylation of neighboring DNA [13].

Since KAISO preferentially binds to methylated DNA, and itself enables recruitment of the DNA methylation modifying machinery (including histone deacetylase-containing nuclear corepressor complex (NCoR) that enhances chromatin condensation [68] and de novo DNA methyltransferases DNMT3a/3b [69] it is possible that when REST and KAISO occupy overlapping sites on DNA, they may compete for binding to DNA, or may interfere with each other’s influence on DNA methylation.

Intersection of the genes with assigned REST-ChIP-seq peaks with those affected by REST knockdown uncovered a number of genes that are high confidence primary targets of REST (Figure 7). Among genes upregulated in REST depleted cells we found genes related to signal release, calcium regulated exocytosis and telencephalon glial cell migration as identified with the REACTOME enrichment analysis. Most of these genes showed decreased expression in G2 and G3 gliomas compared to normal brain samples, with the lowest expression in G4 gliomas (Figure 7B). The expression of these genes was negatively correlated with signatures of G1S and G2M phases of the cell cycle, and positively correlated with NPC-like cellular states as defined by [21]. As NPC-like cellular states are enriched in the proneural GBM [20], originally defined as IDH-MUT GBMs or secondary GBMs (in WHO 2021 classification they would be called G4 astrocytomas) [70], our findings suggest that REST plays a role specifically in these malignant gliomas. Intermediate expression of *REST* in G2/G3 gliomas (Figure 1A) may be enough to maintain a higher expression of the genes contributing to the NPC-like state. High *REST* expression in G4 gliomas might be associated with a strong repression of these genes. This could explain why while REST is a negative prognostic factor in G2/G3 gliomas, it is a positive factor in G4.

## Conclusions

In summary, we identified REST targets, gene regulatory networks and putative REST cooperativity with other TFs that differentially control gene expression in IDH-WT and IDH-MUT gliomas. Among REST targets we found genes involved in glial cell differentiation and ECM organization. Knockdown of REST had a different impact on glioma invasion depending on the IDH phenotype, which is connected to DNA hypermethylation phenotype. We demonstrate that REST–mediated gene transcription activation or repression might be differentially modulated by DNA methylation and by cooperation/competition with other transcription factors, such as KAISO (ZBTB33). The DNA methylation of REST activated genes often showed a positive correlation with gene expression, suggesting that hypermethylation phenotype of IDH-MUT may have a strong impact on these genes. Finally, repression of the canonical REST gene targets may play a more significant role in IDH-MUT grade 2/3 gliomas than in G4 gliomas by maintaining NPC-like cellular state properties. Therefore, REST could be considered as a potential factor in the design of targeted glioma therapies.

## Abbreviations

2HG: 2-hydroxyglutarate
5mC: 5-methylcytosine
α-KG: α-ketoglutarate
AC-like: astrocyte-like
BMP: bone morphogenetic protein
ChIP-seq: chromatin immunoprecipitation-sequencing
coREST: corepressor REST
CpG: cytosine phosphate guanine
dDEG: decreased DEG (a gene differentially expressed in IDH-WT and IDH-MUT that had a lower fold change in siREST-transfected cells than in siCTRL transfected cells)
DEG: differentially expressed gene
ECM: extracellular matrix
ENCODE: The Encyclopedia of DNA Elements
GBM: glioblastoma,WHO grade IV
G2: WHO grade 2
G3: WHO grade 3
G4: WHO grade 4
GO: Gene Ontology
GSC: glioma stem cell
HOCOMOCO: HOmo sapiens COmprehensive MOdel COllection
iDEG: increased DEG (genes differentially expressed in IDH-WT and IDH-MUT that had a higher fold change in siREST-transfected cells than in siCTRL transfected cells)
IDH (1/2): isocitrate dehydrogenase (1 or 2)
IDH-MUT: mutated isocitrate dehydrogenase 1 or 2
IDH-WT: IDH wild type samples (neither IDH1 nor IDH2 were mutated)
LGG: lower grade gliomas, WHO grades 2 and 3
MES-like: mesenchymal-like
NB: normal brain
NPC: neural progenitor cell
NSC: neuronal stem cells
OPC: oligodendrocyte-progenitor cell
PNET: primitive neuroepithelial tumors
PWM: position weighted matrix
RE1: repressor element-1
REST: repressor element-1-silencing transcription factor / RE1 silencing transcription factor/neural-restrictive silencing factor (also known as neuron-restrictive silencer factor NRSF)
REST4: repressor element-1 silencing transcription factor-4
siRNA: short interfering RNA
TCGA: The Cancer Genome Atlas
ZBTB33: Zinc Finger and BTB Domain Containing 33 transcription factor (also known as KAISO)

## Declarations

### Ethics approval and consent to participate

Human tissue methylome and transcriptome data used in this study was derived from publicly available datasets (TCGA, glioma Atlas).

### Consent for publication

Not applicable

### Availability of data and materials

The datasets generated and/or analyzed during the current study are available in the NCBI Gene Expression Omnibus repository (https://www.ncbi.nlm.nih.gov/geo/query/acc.cgi?acc=GSE174308, REST transcription factor holds the balance between the invasion and cell differentiation in IDH-mutant and IDH-wild type gliomas) and in European Genome-Phenome Archive (EGAD00001008986 - Glioma specimens of both IDH-MUT and IDH-WT REST ChIPseq data).

### Competing interests

None declared

### Funding

Studies were supported by the Foundation for Polish Science TEAM-TECH Core Facility project “NGS platform for comprehensive diagnostics and personalized therapy in neuro-oncology”, NCN Symfonia 3 grant nr DEC-2015/16/W/NZ2/00314 and NCN Miniatura-4 grant nr 2020/04/X/NZ2/02126. The use of CePT infrastructure, financed by the European Union, The European Regional Development Fund within the Operational Program “Innovative economy” for 2007–2013, is highly appreciated.

### Authors contributions

MP, BG, KS performed wet lab experiments and sequencing,

MP, BW, AJC, MJD, MJ, MDR analyzed data

MP, BW, AJC, MJ, MJD made the figures

BW, MP, BK, MJD designed the research

BW, MP, MJ, MJD, AJC, MDR, BK wrote the paper.

All the authors read and approved the final manuscript.

## Acknowledgments

We would like to thank Beata Kaza, Sylwia Katarzyna Król, Natalia Ochocka, Iwona Ciechomska, Kamil Wojnicki, Katarzyna Poleszak, Paulina Szadkowska and Aleksandra Ellert-Miklaszewska for their help in the laboratory work and helpful comments and suggestions. We would like to thank Anna Malik and Beata Konikowska for their revision of the manuscript draft.

## Additional material

### File name: Additional file 1

**File format: .pdf**

**Title of data: Differences in gene expression between U87 IDH-MUT and IDH-WT. Description of data: (A)** Volcano plot U87 IDH-MUT vs IDH-WT gene expression comparison. Genes on the right-hand side of the plot have higher expression in IDH-MUT compared to IDH-WT, while genes on the left-hand of the plot have higher expression in IDH-WT compared to IDH-MUT. **(B)** REACTOME pathways analysis of genes expressed differentially between U87 IDH-MUT and IDH-WT glioma cell lines. **(C)** REACTOME pathway analysis of the genes upregulated in U87 IDH-MUT vs IDH-WT cells and TCGA IDH-MUT G2/G3 as compared to IDH-WT gliomas. **(D)** REACTOME pathway analysis of the genes downregulated in IDH-MUT vs IDH-WT U87 cells and TCGA IDH-MUT G2/G3 as compared to IDH-WT gliomas.

### File name: Additional file 2

**File format: .pdf**

**Title of data: Comparison of IDH-MUT in U87 cell lines and primary human grade IV gliomas**

**Description of data: (A)** Sorted values of log2 fold change (log2 FC) for the genes coming from the IDH-MUT vs IDH-WT comparison in U87 cell lines were presented as red dots.

Values for the same genes coming from IDH-MUT vs IDH-WT comparison in primary cell lines were overlaid as gray dots. Percent of log2 FC direction concordance between IDH-MUT vs IDH-WT in U87 cell lines and primary cell lines was calculated. Number of differentially expressed genes is indicated. Black vertical line separates genes expressed higher in IDH-MUT U87 compared to IDH-WT U87 from the genes expressed higher in IDH-WT U87 compared to IDH-MUT U87. **(B)** Genes assigned to extracellular matrix (ECM) organization in Fig3 and downregulated in U87 IDH-MUT vs IDH-WT were presented as zscore heatmap. **(C)** The same genes were visualized using in-house primary IDH-MUT/WT cell lines. In case, when gene was significantly downregulated an asterisk was appended to its name (*=p.adjusted<0.05, **=p.adjusted<0.01, ***=p.adjusted<0.001). **(D)** Bootstrapping result, where significance of obtaining by chance 13 out of 18 significantly downregulated genes was evaluated.

### File name: Additional file 3

**File format:**.**pdf**

**Title of data: Correlation between REST and ECM genes expression levels in TCGA gliomas**

**Description of data:**

Heatmap of the transcript levels of the genes related to extracellular matrix organization GO biological pathway shown in Figure 4C (genes expressed differentially in IDH WT and MUT glioma and modulated by siREST). The columns (patients) were sorted according to decreasing REST gene expression in the tumor sample; left hand side heatmap shows GBM dataset, middle shows LGG IDH-MUT and right-hand side heatmaps shows LGG IDH-WT samples. Asterisk (*) mark genes that had significant correlation of expression with *REST* gene expression in TCGA glioma dataset.

### File name: Additional file 4

**File format: .pdf**

**Title of data: Analysis of genes assigned to REST-ChIPseq peaks specific for IDH-MUT or IDH-WT U87 cells.**

**Description of data: (A)** Differential REACTOME analysis of genes annotated uniquely to either IDH-MUT or IDH-WT REST-ChIPseq peaks. U87 IDH-MUT or IDH-WT promoters (+/- 3kB from transcription start site - TSS) peaks were an input to ChIPseeker analysis with compareCluster function. **(B)** Venn diagram showing the number of TF motifs found in the REST-ChIPseq peaks located within gene promoters in only IDH-WT (red), only IDH-MUT (purple) or both in IDH-WT and IDH-MUT (yellow). **(C)** EnrichR results presenting which TFs based on the ENCODE data were found at genomic positions where REST-ChIPseq peaks were identified in U87 IDH-MUT and U87 IDH-WT. The top panel (yellow) presents results for the peaks common to both IDH-WT and IDH-MUT, the middle (red) - for IDH-WT specific REST-ChIPseq peaks, and the bottom one (purple) for IDH-MUT specific REST-ChIPseq peaks. **(D)** REST and KAISO (ZBTB33) motif occurrence within the gene promoters bound in REST-ChIPseq. Venn diagrams show the number of gene promoter sequences that contained REST or KAISO motifs and were (I) common between IDH-WT and IDH-MUT, (II) unique to IDH-WT, and (III) unique to IDH-MUT.

### File name: Additional file 5

**File format: .pdf**

**Title of data: DNA methylation pattern of the REST-ChIPseq peaks (summit ± 100bp) in tumors (glioma Atlas).**

**Description of data:** REST ChIP-seq peaks **(A)** common, **(B)** IDH-MUT-specific or **(C)** IDH-WT-specific that were differentially methylated in the following pairs of glioma tumor types; **(D-E)** Mean REST-ChIPseq peak DNA methylation of the peaks differentially methylated between G2/G3 IDH-WT and G2/G3/G4 IDH-MUT glioma samples. Heatmaps are arranged in order of decreasing difference in methylation between IDH-WT and IDH-MUT G2/G3/G4 glioma samples; **(D)** Peaks common for IDH-WT and IDH-MUT (n=128); **(E)** REST ChIP-seq peaks unique to IDH-MUT (n=4); **(F)** REST ChIP-seq peaks unique to IDH-WT (n=32).

### File name: Additional file 6

**File format: .pdf**

**Title of data: Characterization of ChIPseq peaks and TF motifs in REST-repressed and REST-activated genes.**

**Description of data: (A)** Gene Ontology Biological Process gene enrichment analysis for repressed REST targets (left panel) and activated REST targets (right panel). **(B)** Clustering of TF motifs characteristic for genes activated by REST, repressed by REST, and common to both REST-activated and REST-repressed, based on TF PWMs. **(C-D)** Distribution of KAISO motifs within the promoters of the genes activated **(C)** and repressed **(D)** by REST.

**(A)** Distribution of REST motifs within the promoters of the genes repressed by REST. **(F)** DNA sequence logos of the REST binding motifs PWMs; **(G)** DNA sequence logos of the KAISO binding motifs PWMs.

### File name: Additional file 7

**File format:.doc**

**Title of data: Additional Table 1**

**Description of data:** Number of cytosines within REST or KAISO motifs within REST-ChIPseq peaks assigned to REST-repressed or -activated targets. The peaks were categorized according to whether they had detected binding motifs of both REST and KAISO, or just one of them.

### File name: Additional file 8

**File format: .doc**

**Title of data: Additional Table 2**.

**Description of data:** Number of sites where individual REST or KAISO motifs were detected within REST-ChIPseq peaks derived from the cell lines. Sites discovered as differentially methylated among glioma tumor groups were matched to individual REST or KAISO motifs and then to peaks that were paired with REST-repressed or -activated targets. The number of all detected sites of REST or KAISO motifs and the number of differentially methylated sites were significantly different (X-squared = 34.763, df = 14, p-value = 0.001593). Asterisk indicates differences in DNA methylation larger than expected by chance (*). ND - no data.

### File name: Additional file 9

**File format: .doc**

**Title of data: Additional Table 3**.

### File name: Additional file 10

**File format: .pdf**

**Title of data: REST-ChIP-seq peaks DNA methylation level, of U87 IDH-WT and U87 IDH-MUT cell lines samples treated with siCTRL and siREST, in a resolution of single cytosine loci.**

**Description of data:** List of REST repressed or REST activated genes containing differentially methylated sites in REST or KAISO motifs within the associated REST-ChiPseq peaks. A) Difference versus log10 fold change (FC) in DNA methylation between siREST and siCTRL samples of U87 IDH-WT cell line; B) as in A but for U87-MUT cell line; C) Difference versus fold change in DNA methylation between siCTRL IDH-MUT and siCTRL WT samples; D) as in C but for siREST IDH-MUT and siREST IDH-WT samples; In A-D: light gray - no difference, red - decreased methylation (difference <= -0.15), green - increased methylation (difference > =0.15); E) Difference in cytosine methylation between U87-MUT and U87-WT cell lines in siREST and siCTRL samples; F) Cytosines within REST-ChIPseq peaks were divided into three categories (WT-specific, MUT-specific, common) and presented as percentage of single cytosines that overlapped with one of the three types of REST-ChIPseq peaks. “Background” refers to all cytosines within REST-peaks, “Differential” represents non-light gray cytosines shown C&D, “Higher” (red) - cytosines with higher DNA methylation in IDH-MUT, Lower (green) - cytosines with lower DNA methylation in IDH-MUT; G) log10 fold change between percentage of “Differential” cytosines to “Background” cytosines in each of the three REST-ChIPseq peaks categories: common, WT-, MUT-specific.

### File name: Additional file 11

**File format: .pdf**

**Title of data: GO Biological Pathway enrichment for the genes identified in REST-ChIPseq and and REST knockdown experiments.**

**Description of data: (A)** GO BP analysis for the genes that were upregulated or **(B)** downregulated by siREST and targeted by REST in the ChIPseq experiment. ChIPseq experiments on both U87 IDH-MUT, IDH-WT cell lines and from freshly resected human tumors show a good overlap of the number of genes annotated to ChIPseq peaks.

## Notes

https://github.com/mdraminski/transcripionFactorREST

https://www.ncbi.nlm.nih.gov/geo/query/acc.cgi?acc=GSE174308

### Competing Interest Statement

The authors have declared no competing interest.

### Summary of Updates

To provide a broader background for our findings, we included results from additional 12 human primary grade 4 glioma cell lines, ten of which were classified as GBM IDH-WT and two as grade 4 astrocytoma IDH-MUT, and compared them to U87 cell lines used in our analysis. The resulting data is presented in Additional File 2. We have also added data on DNA methylation within the REST-ChIPseq peaks from U87 IDH-WT and U87 IDH-MUTcells with REST knockdown (presented in Additional File 10). We added some changes to the manuscript text, which include: - Changes to the numerals describing glioma grades form Roman to Arabic according to WHO 2021 terminology and update of the term "secondary glioblastomas" to "astrocytoma grade 2, 3, 4"; - A short fragment on E2F involvement in glioma progression was added in reference to Figure 5 E; - An additional paragraph addressing the possible KAISO / REST competition in the context of DNA methylation was added to the discussion; - A comment on the possibility of different REST contributions to different cell migration types was added to the discussion; - A paragraph addressing what is known about REST in glial versus neuronal differentiation has been extended in the discussion.

