## Supplementary figures and images for "Comprehensive analysis of the transcription factor REST regulatory networks in IDH-mutant and IDH-wild type glioma cells and gliomas"

### Additional File 1

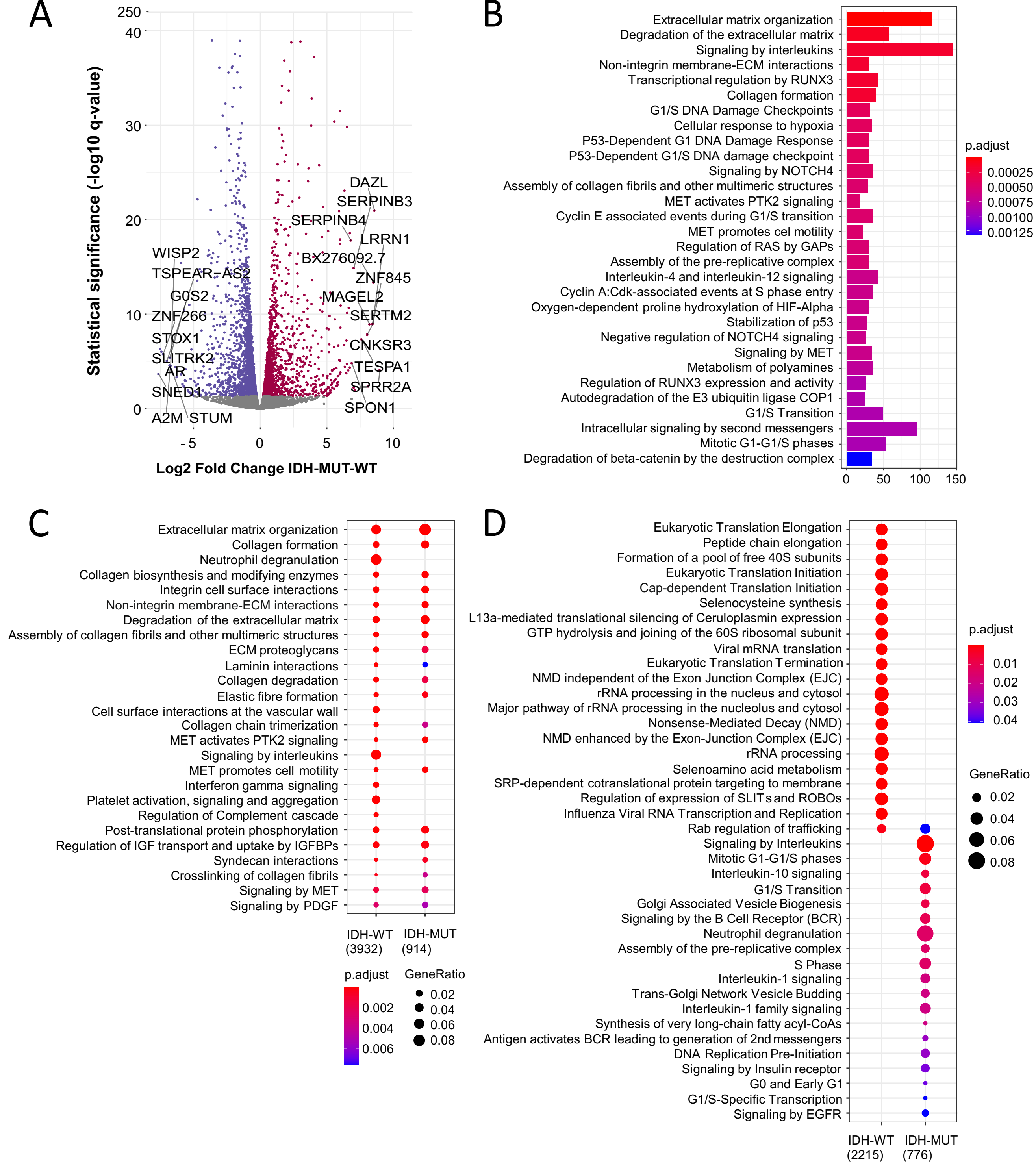

### Additional File 2

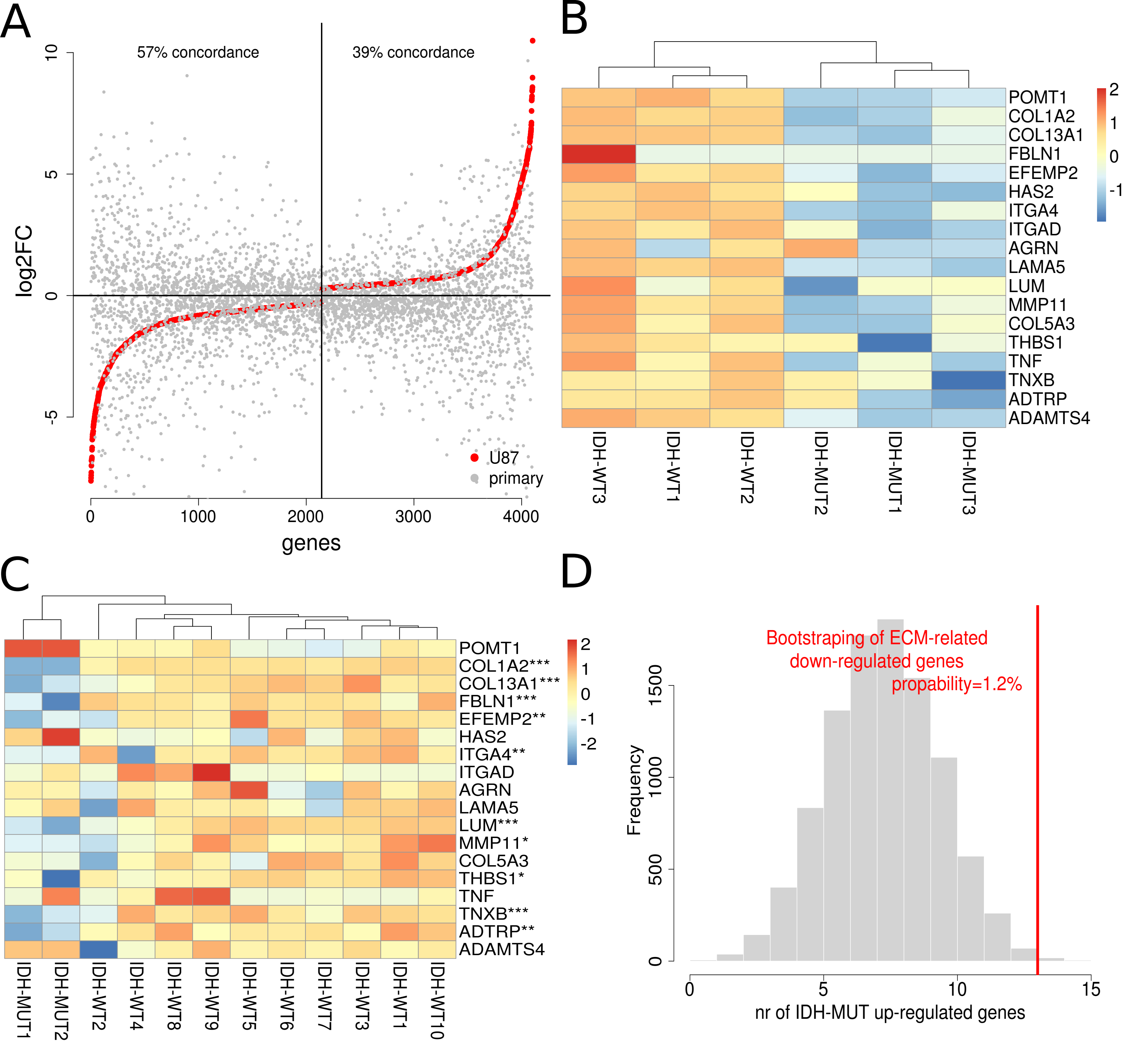

### Additional File 3

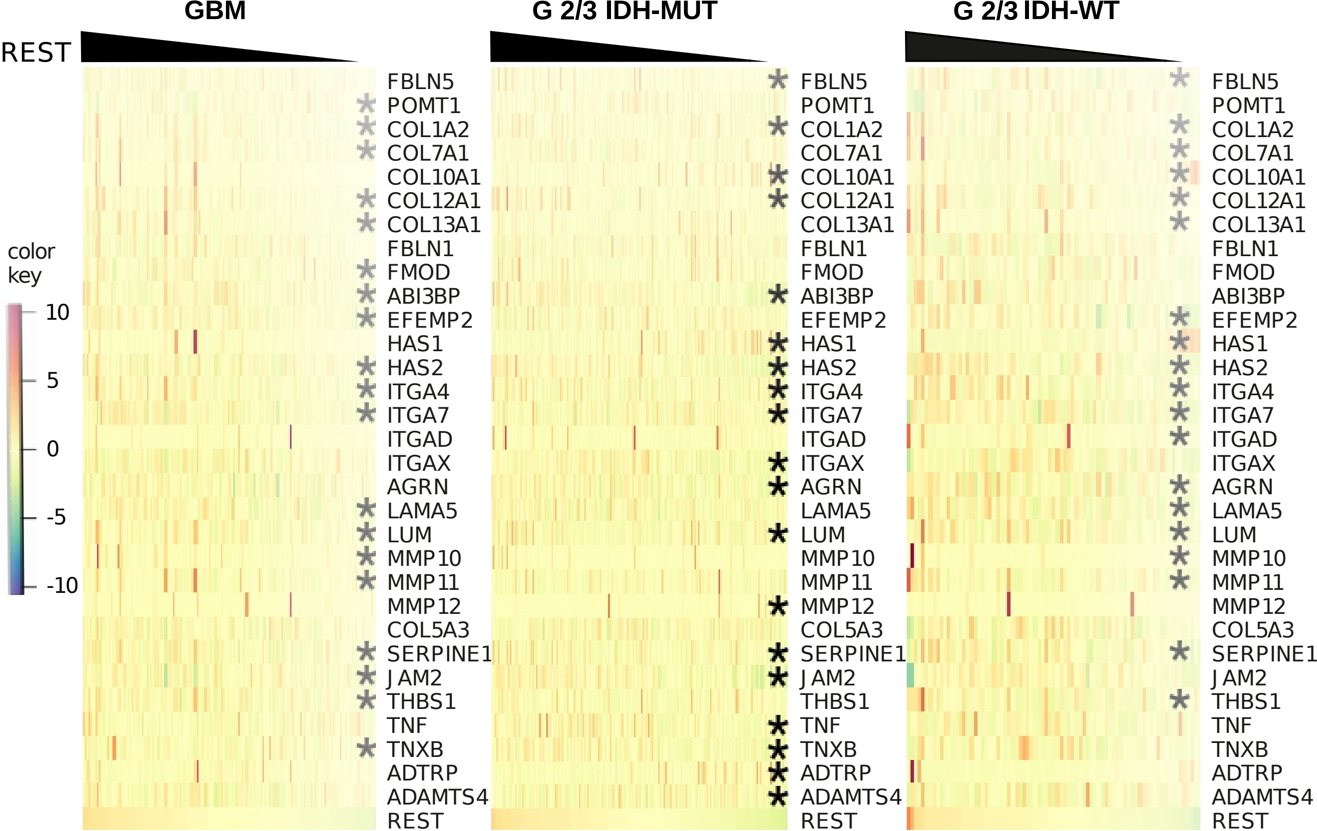

### Additional File 4

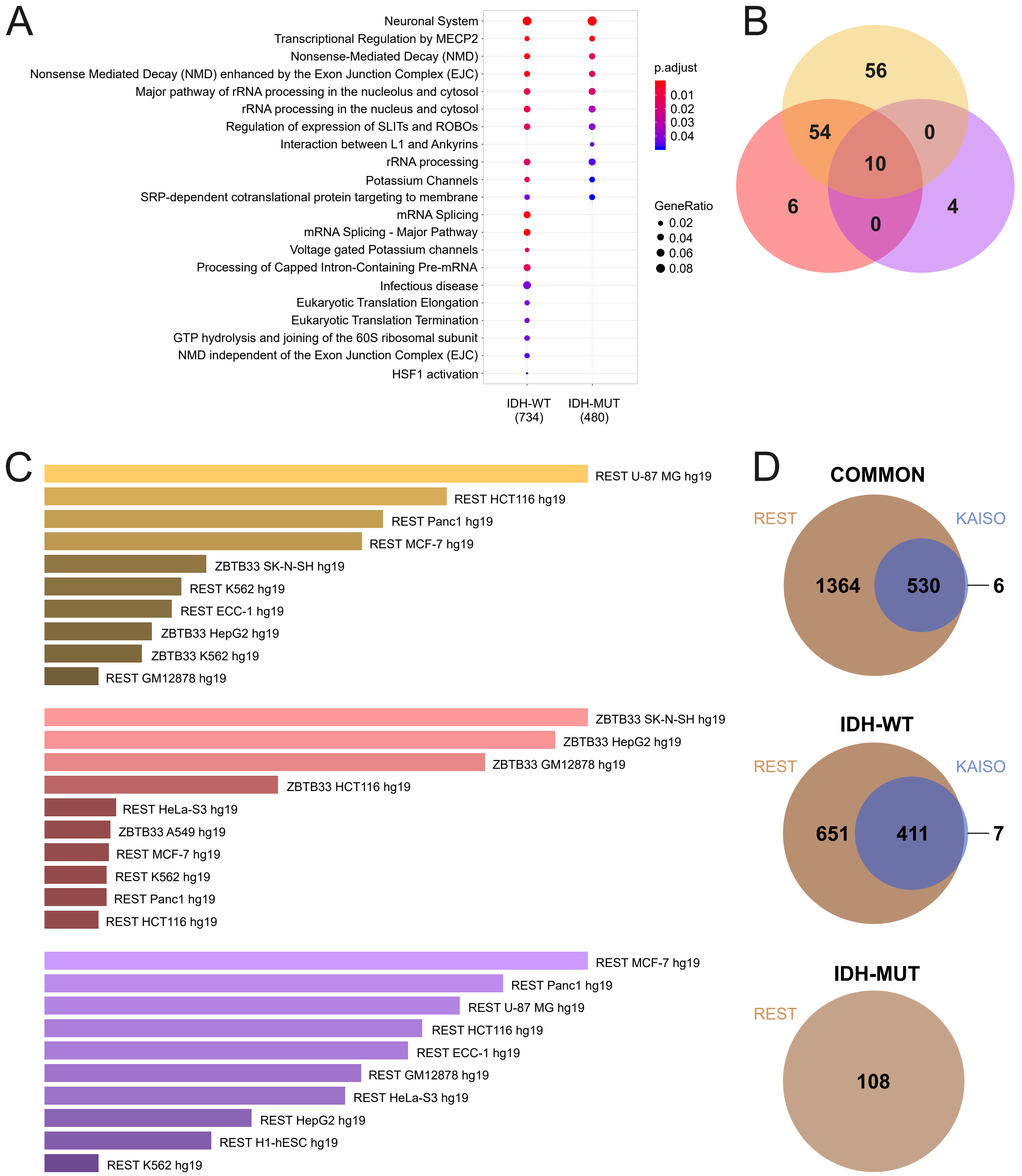

### Additional File 5

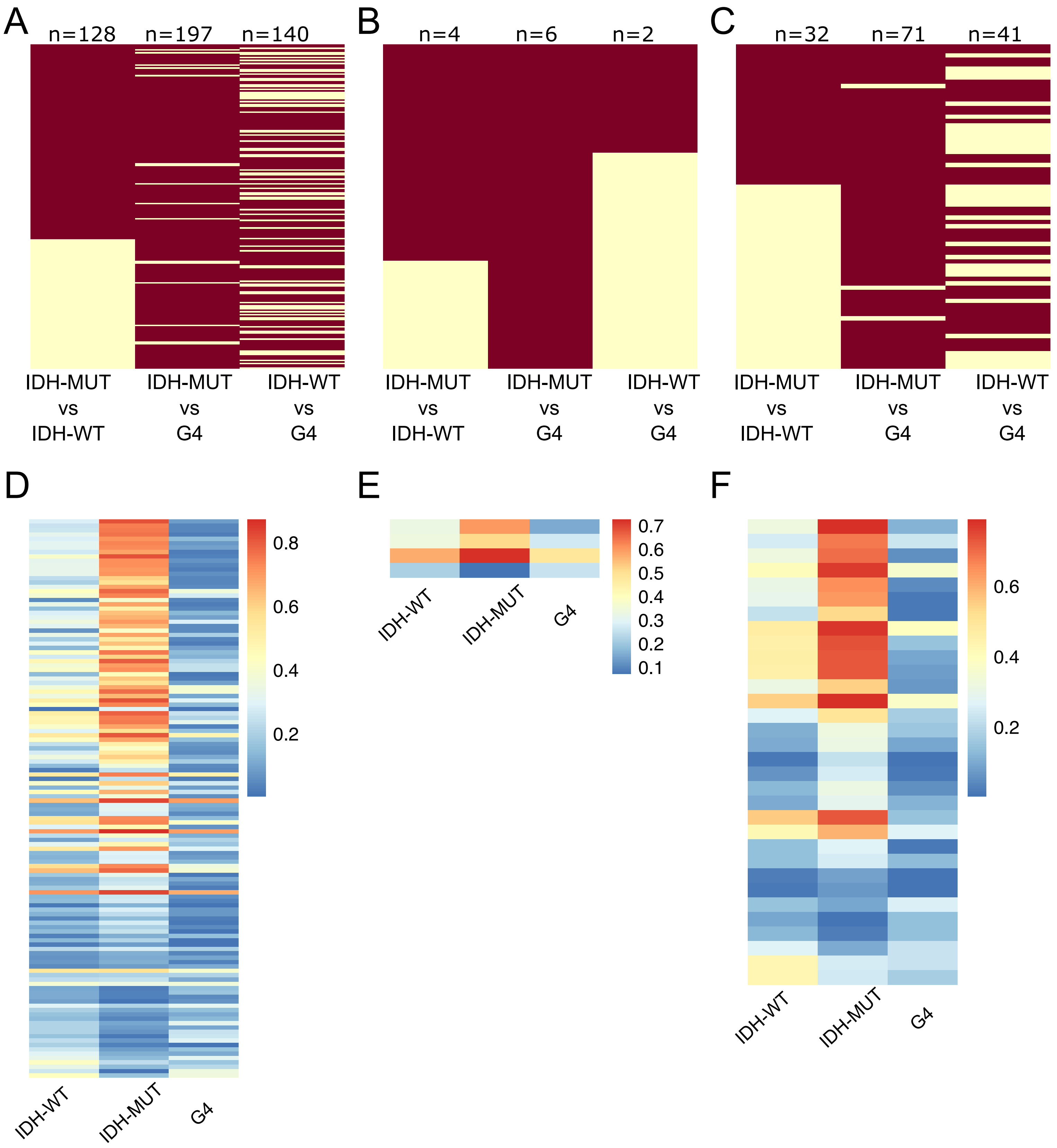

### Additional File 6

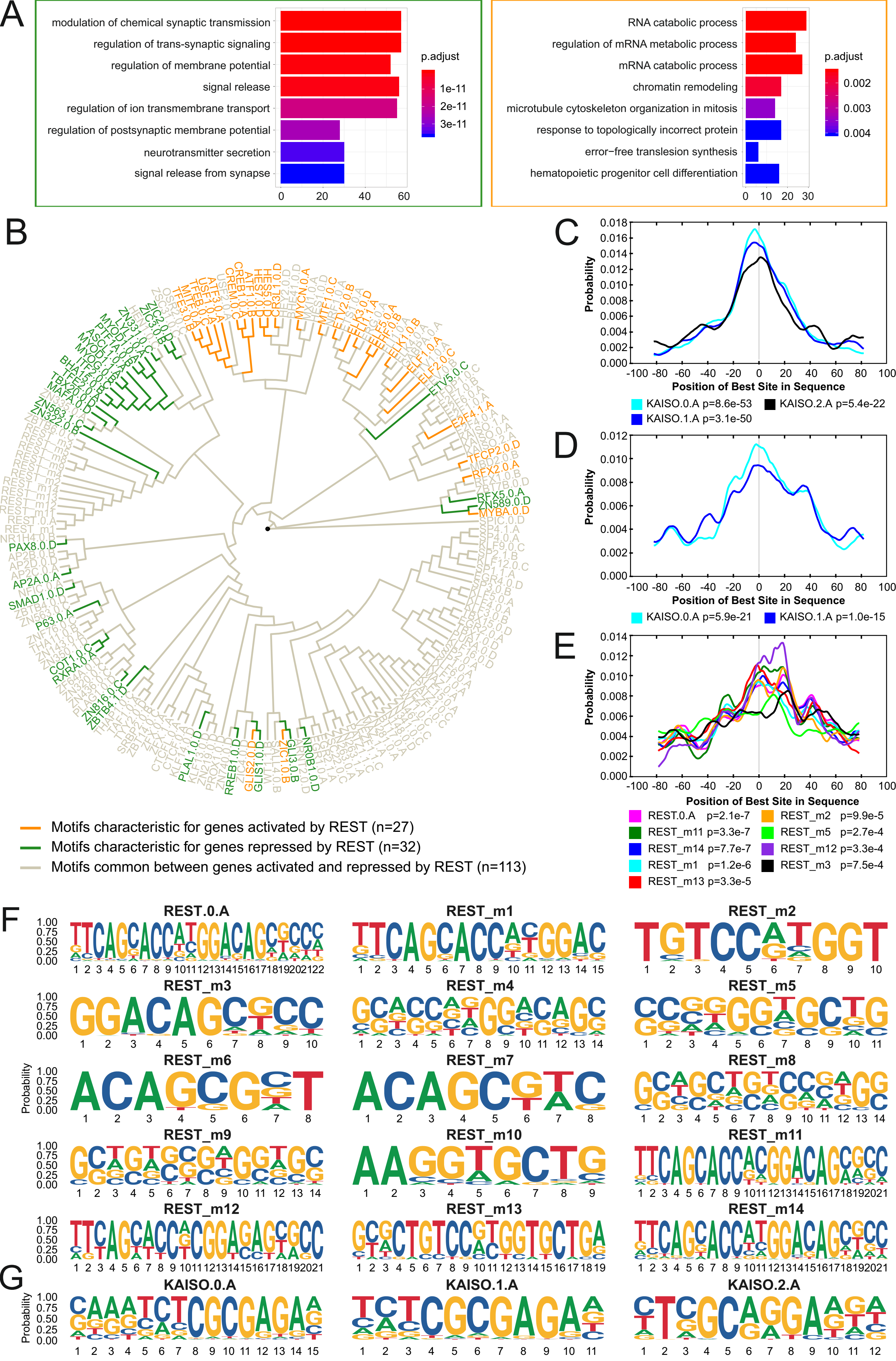

### Additional File 10

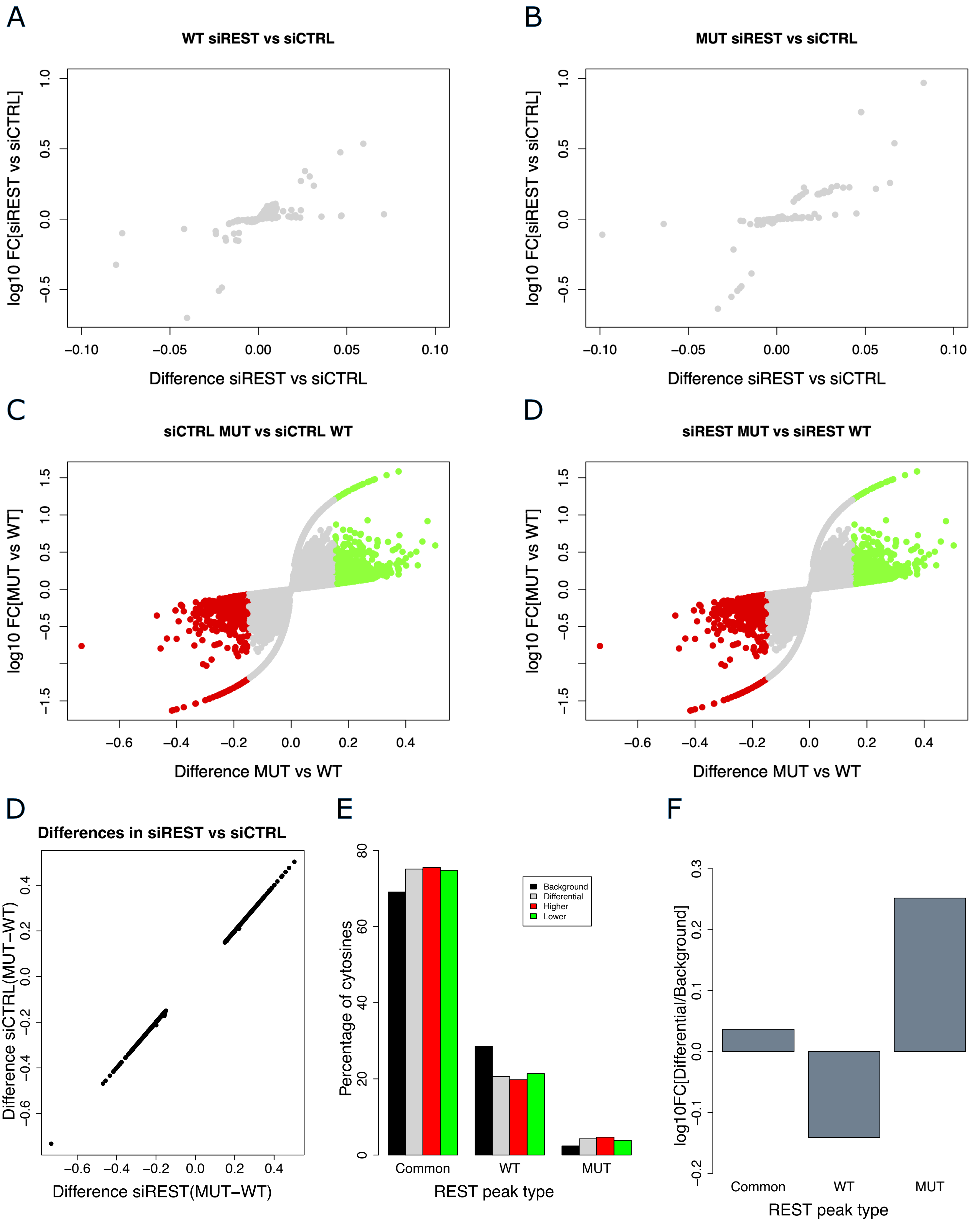

### Additional File 11

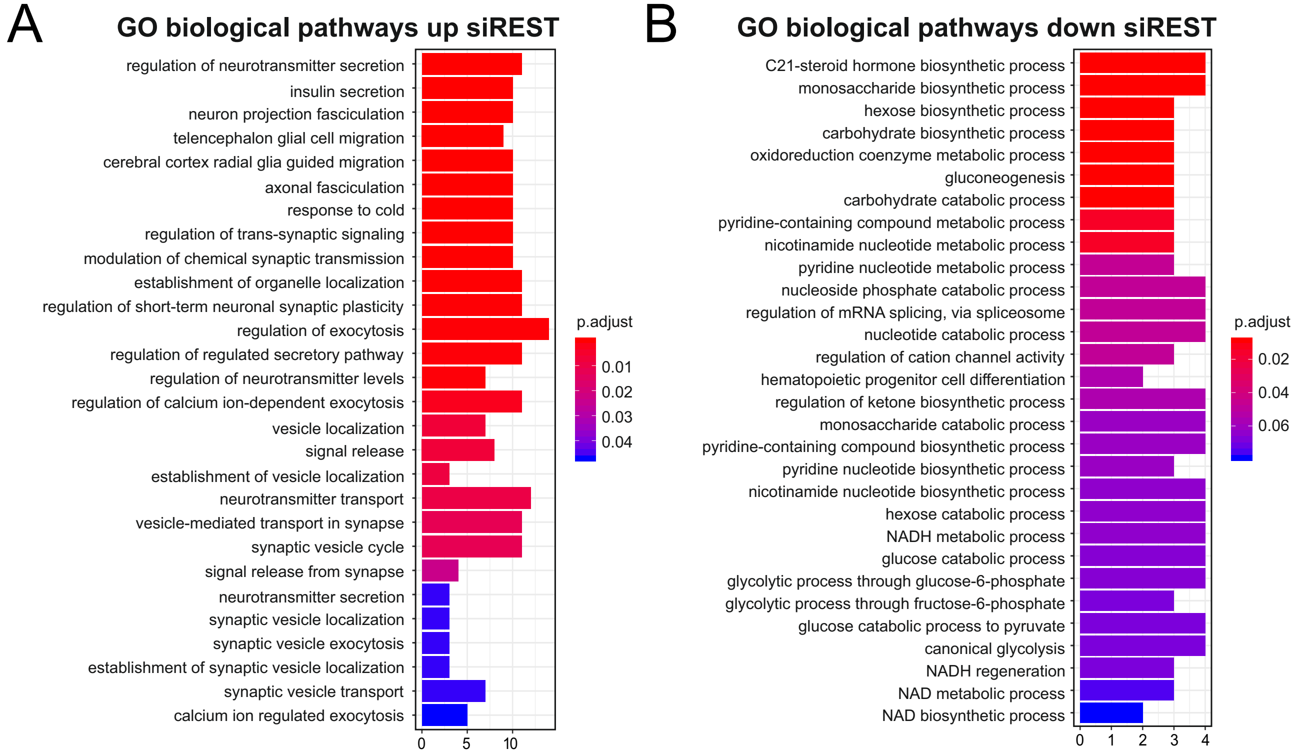
